# The GxcM-Fbp17/RacC-WASP signaling cascade regulates polarized cortex assembly in migrating cells

**DOI:** 10.1101/2022.11.14.515780

**Authors:** Dong Li, Yihong Yang, Yingjie Wang, Xiaoting Chao, Jiafeng Huang, Shashi P. Singh, Chengyu Zhang, Jizhong Lou, Pu Gao, Shanjin Huang, Huaqing Cai

**Author notes:** These authors contributed equally to this work (Dong Li, Yihong Yang).

## Abstract

The actin-rich cortex plays a fundamental role in many cellular processes. Its architecture and molecular composition vary across cell types and physiological states. The full complement of actin assembly factors driving cortex formation and how their activities are spatiotemporally regulated remain to be fully elucidated. Using *Dictyostelium* as a model for polarized and rapidly migrating cells, we show that GxcM, a RhoGEF localized specifically in the rear of migrating cells, functions together with F-BAR protein Fbp17, a small GTPase RacC, and the actin nucleation-promoting factor WASP to coordinately promote Arp2/3 complex-mediated cortical actin assembly. Over-activation of this signaling cascade leads to excessive actin polymerization in the rear cortex, whereas its disruption causes defects in cortical integrity and function. Therefore, different from its well-defined role in the formation of the front protrusions, the Arp2/3 complex-based actin carries out a previously unappreciated function in building the rear cortical subcompartment in rapidly migrating cells.

## Introduction

The cell cortex is defined as a thin layer of filamentous actin, myosin motors, and regulatory proteins beneath the plasma membrane (Svitkina, 2020). Assembly and contraction of this actin meshwork generates cortical tension, which enables cells to resist external mechanical stresses, change shape, and exert forces (Chugh et al., 2017; Kelkar et al., 2020). Consequently, the cortex plays a critical role in a variety of cellular processes, including division, migration, and morphogenesis (Heisenberg and Bellaiche, 2013; Salbreux et al., 2012). The mechanical properties of the cortex are key to its physiological function. Changes in cortical mechanics can originate from changes in the architecture of the actin network, which, as a viscoelastic structure, can be remodeled on a time scale of seconds (Fritzsche et al., 2016; Koenderink and Paluch, 2018; Svitkina, 2020). Rapid actin rearrangements may enable cells to modify their shapes for rapid adaptation to changes in the extracellular environment. However, the complete inventory of assembly factors driving formation of the actin cortex and how their activities are spatiotemporally controlled are not well understood.

Actin polymerization is mainly initiated by two classes of nucleators: the Arp2/3 complex, which creates branches at the sides of preexisting filaments to generate a dense meshwork, and formins, which nucleate and elongate long and linear actin filaments (Pollard, 2007). Recent biochemical and functional studies have implied that both Arp2/3 and formins are involved in the formation of the cortical actin cytoskeleton, though their relative contributions vary among cell types (Bovellan et al., 2014; Chan et al., 2019; Chugh et al., 2017; Fritzsche et al., 2016; Litschko et al., 2019; Rosa et al., 2015; Severson et al., 2002). For example, the Arp2/3 complex is largely dispensable for formation of the cell cortex of mitotic epithelial cells within the fly notum and early *Caenorhabditis elegans* embryos (Rosa et al., 2015; Severson et al., 2002). However, in M2 melanoma cells and mitotic HeLa cells, both Arp2/3 and the Diaphanous-related formin (DRF) mDia1 contribute to cortical F-actin, though with different effects on cortical integrity and cell behavior. Although depletion of mDia1 arrests cell division, depletion or perturbation of Arp2/3 complex alone does not. Intriguingly, Arp2/3 complex inhibition potentiates the effect of mDia1 depletion, suggesting synergistic activities of the two types of actin nucleators (Bovellan et al., 2014).

The activities of Arp2/3 complex and DRFs are tightly regulated. The DRFs adopt an autoinhibited conformation through intramolecular interactions; they can be activated by binding with Rho or Ras family GTPases (Breitsprecher and Goode, 2013; Pollard, 2007). The activation of Arp2/3 complex relies on nucleation-promoting factors (NPFs) of the Wiskott-Aldrich syndrome protein (WASP) family, which consists of two principal classes of protein: WASPs and WASP family verprolin homologous (WAVE) complex (also called SCAR complex). Both of them contain a C-terminal VCA domain that binds and activates Arp2/3 in response to numerous inputs, including Rho GTPases, phosphoinositide lipids, SH3 domain-containing proteins, kinases, and phosphatases (Pollard, 2007; Pollitt and Insall, 2009). Recently, the presence of several Arp2/3 NPFs in the cortex has been shown by proteomic analysis of cortex-containing blebs and fluorescent microscope imaging (Bovellan et al., 2014; Cao et al., 2020). Modulating the activities of Arp2/3 and formins is thought to fine-tune the composition, structural organization, and mechanics of the actin cortex.

*Dictyostelium discoideum* provides a valuable system for studying the architecture of the actin cortex in rapidly moving and polarized cells. *Dictyostelium* cells exemplify amoeboid migration, which is characterized by weak adhesions, actin-rich protrusions or blebs in the front, and actomyosin-driven contraction in the rear (Devreotes et al., 2017; Lammermann and Sixt, 2009; Paluch et al., 2016). Previous studies have demonstrated that three DRFs (ForA, ForE, and ForH), which are recruited and activated by the RhoA-like GTPase RacE, act together to safeguard cortical integrity in the rear of migrating cells (Litschko et al., 2019). Some evidence suggests that the Arp2/3 complex also takes part in building the rear cortex. First, cryo-electron tomography of peripheral regions in *Dictyostelium* cells revealed isotropic actin-filament arrays (Medalia et al., 2002). Second, a significant amount of cortical actin meshwork remained in cells in which all three DRFs were deleted (Litschko et al., 2019). Third, antibodies to the Arp2/3 complex stained around the cortex in related amoeba cells (Machesky et al., 1994; Mullins et al., 1997). However, as prominent Arp2/3 activities are mainly found at the leading-edge of cells, including the structures of pseudopods and macropinocytic cups (Davidson et al., 2018; Veltman et al., 2012; Veltman et al., 2016; Yang et al., 2021), whether and how the Arp2/3 complex can be brought to the rear to promote cortical actin generation remains unknown.

In this study, we show that GxcM, a RhoGEF protein localized in the rear of *Dictyostelium* cells, functions together with F-BAR protein Fbp17, a Rac family GTPase RacC, WASP, and Arp2/3 complex in a signaling cascade to coordinately regulate the formation of a rear cortical actin subcompartment and maintain cortical integrity.

## Results

### RhoGEF protein GxcM localizes in the rear of migrating cells

We reasoned that, for Arp2/3-based actin to be involved in formation of the rear cortex, asymmetric positioning of upstream regulators may be required. Rho family GTPases and their respective guanine nucleotide exchange factors (GEFs) are well-known regulators of Arp2/3-dependent actin polymerization. Intriguingly, in a previous screen aiming to identify pleckstrin homology (PH) domain-containing proteins that localize in the rear of cells (Li et al., 2022), we found a putative RhoGEF named GxcM (Fig. 1A). In randomly moving vegetative cells, GxcM-RFP and the well-characterized leading-edge marker PHcrac-GFP (Parent et al., 1998; Yang et al., 2021), a sensor for PIP_3_/PI(3,4)P_2_, exhibited an inverse distribution, the latter accumulating in regions where GxcM was depleted (Fig. 1B). In cells migrating under agarose along a folic acid gradient, GxcM-GFP localized more strongly to the side and rear of cells, resulting in an apparent rear-to-front gradient in its plasma membrane association (Fig. 1H; Video 1). Furthermore, as previously reported for rear proteins (Iijima and Devreotes, 2002; Li et al., 2022; Swaney et al., 2015), GxcM-GFP transiently translocated from the membrane to the cytosol upon global chemoattractant stimulation, and the translocation did not require an intact actin cytoskeleton (Fig. 1C; Fig. S1A).

**Figure 1.**
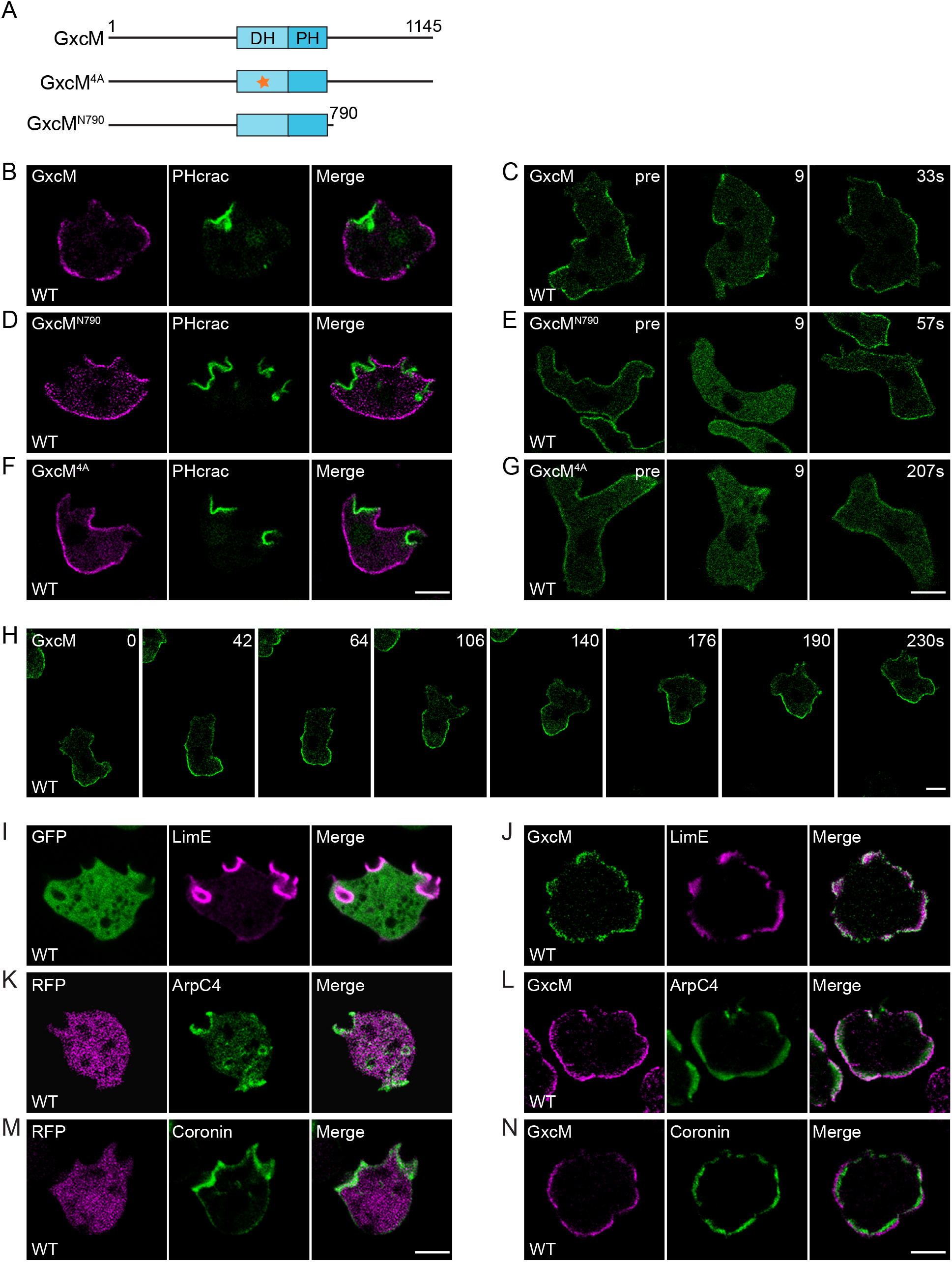
GxcM localizes in the rear of *Dictyostelium* cells. (A) Schematic representation of GxcM, GxcM^4A^, and GxcM^N790^. The star indicates alanine mutations (aa 593-596) in the DH domain. (B, D, F) Inverse distribution of PHcrac-GFP and RFP-tagged GxcM (B), GxcM^N790^ (D), or GxcM^4A^ (F) in randomly migrating cells. (C, E, G) Transient translocation of GFP-tagged GxcM (C), GxcM^N790^ (E), and GxcM^4A^ (G) in response to cAMP stimulation added at time 0 (pre: before stimulation). (H) Time-lapse imaging of GxcM-GFP in cells migrating under agarose along a folic acid gradient. (I-J) Localization of RFP-LimEΔcoil and the GFP control (I) or GxcM-GFP (J) in randomly migrating cells. (K-L) Localization of GFP-ArpC4 and the RFP control (K) or GxcM-RFP (L) in randomly migrating cells. (M-N) Localization of GFP-Coronin and the RFP control (M) or GxcM-RFP (N) in randomly migrating cells. Scale bars, 5 μm.

We generated truncations to investigate the localization requirement of GxcM. GxcM contains a RhoGEF domain (also known as Dbl homology domain or DH domain, 455-638 aa) and a PH domain (626-770 aa) in the central region (Fig. 1A). Deleting the C-terminus of GxcM (GxcM^N790^) did not change its reciprocal localization to PHcrac or response to stimulation (Fig. 1, D and E; Fig. S1B), whereas further deletion by removing the DH and PH domains (GxcM^N455^) caused it to completely dissociate from the membrane (Fig. S1C). The DH and PH domains, though necessary, were not sufficient to target GxcM because the truncation containing only the two domains (GxcM^455-770^) localized in the cytoplasm (Fig. S1D). Sequence alignment of GxcM with human ECT2, a RhoGEF involved in the regulation of cytokinesis (Tatsumoto et al., 1999), revealed four conserved residues (PVQR, 593-596 aa) within the DH domain (Fig. S1F). This stretch of residues has been shown to be essential for the GEF activity of ECT2 (Su et al., 2011). Mutating the four residues to alanines (GxcM^4A^) did not affect the localization or chemoattractant-induced translocation of GxcM (Fig. 1, F and G; Fig. S1E). Therefore, GxcM is a RhoGEF localized selectively in the rear of *Dictyostelium* cells.

### Overexpression of GxcM induces Arp2/3-mediated actin assembly in the rear cortex

To dissect the potential role of GxcM in cortex assembly, we generated *gxcM* knockout (*gxcM* ^−^) cells (Fig. S2A) and examined the cellular activities known to rely on cell shape remodeling and cortex integrity (Litschko et al., 2019; Ramalingam et al., 2015). Development of *gxcM* ^−^ cells on bacterial lawns was indistinguishable from that of wild-type (WT) cells (Fig. S2B). When assayed in shaken suspension, a condition used previously to reveal cytokinesis defects in the formin and RacE mutants (Litschko et al., 2019), *gxcM* ^−^ cells exhibited only a mild cytokinesis defect, with the vast majority of *gxcM* ^−^ cells being mono- or bi-nucleated like WT cells, and only approximately 7.9% exhibited three or more nuclei (Fig. S2C). In random motility assays and under agarose chemotaxis assays, *gxcM* ^−^ cells migrated with a speed and directness comparable to that of WT (Fig. S2, D and E). Thus, disruption of *gxcM* did not appear to markedly impair cell cortex integrity.

Despite the mild phenotypes in *gxcM* ^−^ cells, from the localization experiments (Fig. 1, B and H), we observed that overexpression of GxcM from extrachromosomal vectors seemed to alter cell morphology and behavior. Time-lapse imaging further revealed the impact of GxcM overexpression. Cells expressing Teep1-RFP, a rear protein we characterized previously (Li et al., 2022), GxcM^4A^-RFP, or GxcM^N790^-RFP exhibited a polarized morphology and formed one or two protruding fronts at a given time in the form of pseudopods or macropinocytic cups that were labeled by PHcrac-GFP (Fig. S3A; Video 2). In contrast, GxcM-RFP-expressing cells were less polar and produced rounded GxcM-marked protrusions at peripheral regions depleted of PHcrac (Fig. S3B; Video 2). These GxcM-enriched structures did not effectively cause cell displacement or macropinocytosis, but generated merely outward bulges of the cell boundary. Consequently, GxcM overexpressing cells exhibited reduced motility and macropinocytosis (Fig. S3, C and D). We verified that these phenomena relied on overexpression of GxcM, as expression of GxcM-GFP from a genome-integrated expression cassette, which did not generate signals bright enough for live cell imaging, did not seem to cause morphological change of the cell (Fig. S3E).

To examine whether GxcM overexpression altered cell morphology by mediating changes in the actin cytoskeleton, we co-expressed GxcM-GFP with LimEΔcoil-RFP, a marker for newly polymerized actin (Bretschneider et al., 2004). In cells expressing the GFP control, the bulk of LimEΔcoil signal was found at the pseudopod or macropinocytic cup region, and less prominent signals were seen at the rear and lateral sides (Fig. 1I; Video 3). On the other hand, in GxcM-GFP-overexpressing cells, the LimEΔcoil signal was highly concentrated at peripheral regions enriched with GxcM (Fig. 1J; Video 3), suggesting that GxcM expression may induce ectopic actin polymerization. Further analyses using different actin activity reporters showed that the GxcM-induced actin assemblies were likely Arp2/3-based, because they could be marked by ArpC4, an Arp2/3 complex subunit, and Coronin, a central constituent of the Arp2/3-mediated actin network (Fig. 1, K-N; Video 3). Similar morphological changes and reorganization of actin structures were observed in cells expressing untagged GxcM, ruling out non-specific effects of the engineered fluorescent tag (Fig. S3F).

We also observed the actin activity markers in GxcM-overexpressing cells chemotaxing under agarose along a folic acid gradient. Though LimEΔcoil, AprC4, and Coronin were mainly detected in protrusions at the migrating front in control cells (Fig. 2, A and C; Fig. S4A; Video 4), a significant fraction of these reporters were localized to the rear and lateral sides where intensive GxcM signals were observed in GxcM-overexpressing cells (Fig. 2, B and D; Fig. S4B; Video 4). In addition, traveling actin waves were frequently seen in the rear of cells upon GxcM overexpression. Perhaps as expected, the rearrangement of Arp2/3 activity and F-actin interfered with cell function. Although the GxcM-overexpressing cells were still able to orient and move towards the higher concentration of folic acid, they migrated with considerably slower speed (Fig. S4C), highlighting the need to fine-tune the level of GxcM. Cytokinesis was also weakly impaired in these cells (Fig. S3G). These results suggest that GxcM may function as a cortex assembly factor for promoting Arp2/3-based actin polymerization in the rear of cells, and its overexpression further boosts such activity, resulting in over-assembly of actin and disruption of cell function.

**Figure 2.**
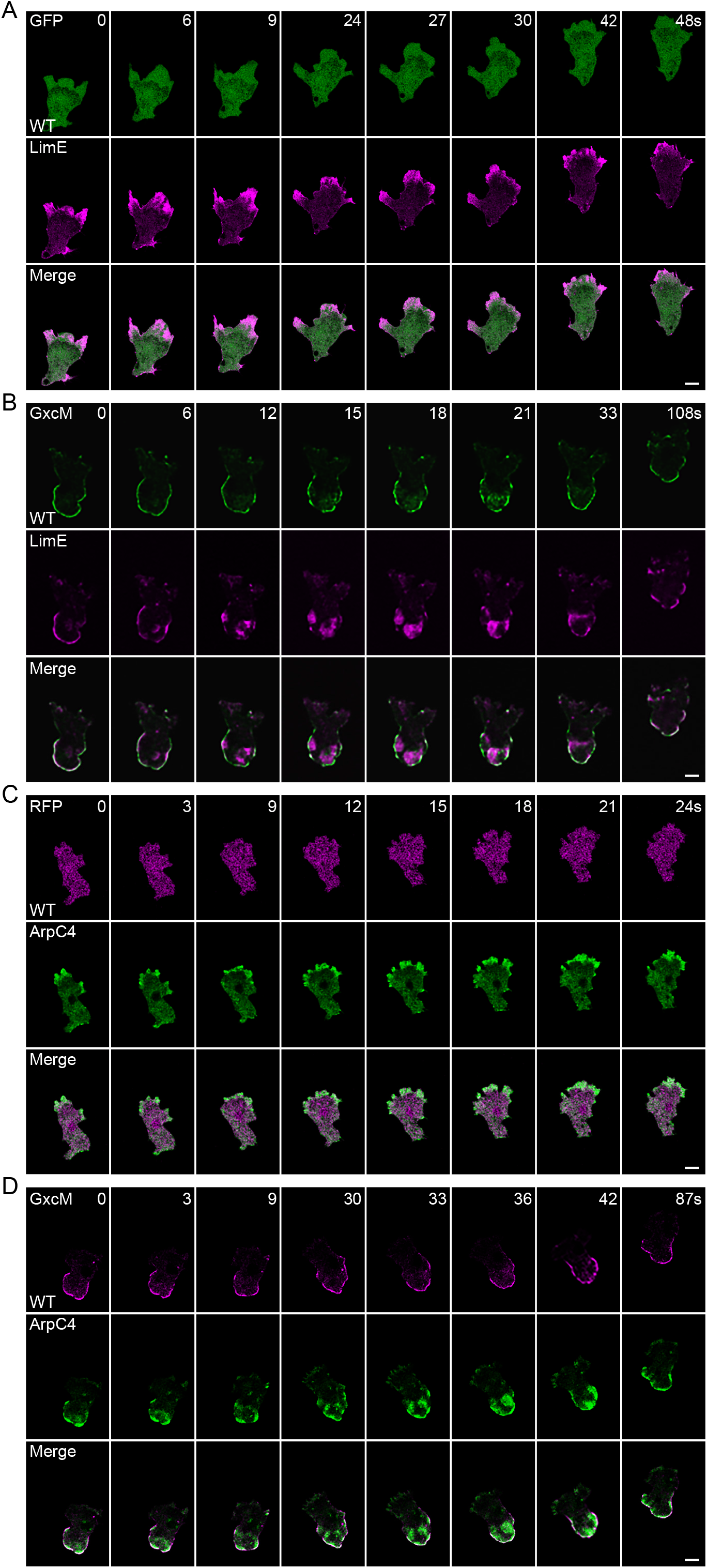
GxcM overexpression induces cortical actin assembly. (A-B) Time-lapse imaging of RFP-LimEΔcoil and GFP (A) or GxcM-GFP (B) in WT cells migrating under agarose along a folic acid gradient. (C-D) Time-lapse imaging of GFP-ArpC4 and RFP (C) or GxcM-RFP (D) in WT cells migrating under agarose along a folic acid gradient. Scale bars, 5 μm.

Overexpression of the GxcM truncations that localized in the cytoplasm, including GxcM^N455^-GFP and GxcM^455-770^-GFP, did not alter the cell morphology or trigger actin polymerization. Overexpression of GxcM^N790^-GFP and GxcM^4A^-GFP also failed to induce actin assembly, as evidenced by the preferential leading-edge enrichment of LimEΔcoil, AprC4, and Coronin in randomly migrating (Fig. S5, A-F) and chemotaxing cells (Fig. S5, G and H). Therefore, the cortical actin assembly-promoting activity of GxcM requires an intact C-terminus, as well as the GEF domain.

### The actin assembly-promoting activity of GxcM relies on interaction with F-BAR protein Fbp17

We noted that the C-terminus (791-1145 aa) of GxcM is remarkably proline-rich (Fig. S6A). As a diverse array of actin regulators contain proline-binding modules, such as SH3 and EVH1 domains (Ball et al., 2005; Holt and Koffer, 2001), we speculated that GxcM may function by association with such proteins via its C-terminus. To identify binding partners of GxcM, we immunoprecipitated GxcM-GFP, GxcM^N790^-GFP, or Teep1-GFP from cell lysates and performed mass spectrometry (MS) analysis. A SH3 domain-containing protein (gene ID, DDB_G0271812) was uniquely enriched in the immunocapture of GxcM-GFP (Fig. S6, B and C). Sequence analysis revealed that this protein is a homolog of the FBP17 family of Fes/CIP4 homology-Bin-Amphiphysin-Rvs (F-BAR) proteins (Fig. S7A). This family includes three closely related proteins, formin-binding protein 17 (FBP17), transactivator of cytoskeletal assembly-1 (TOCA-1), and Cdc42-interacting protein 4 (CIP4), which are all potent activators of Arp2/3-dependent actin polymerization (Chen et al., 2013; Ho et al., 2004; Takano et al., 2008; Tsujita et al., 2006). DDB_G0271812 shares 21.3% identity and 38.6% similarity to human FBP17, and as FBP17, it contains an F-BAR, a homology region 1 (HR1), and a SH3 domain (Fig. 3A; Fig. S7, B and C). Therefore, we named this protein Fbp17.

**Figure 3.**
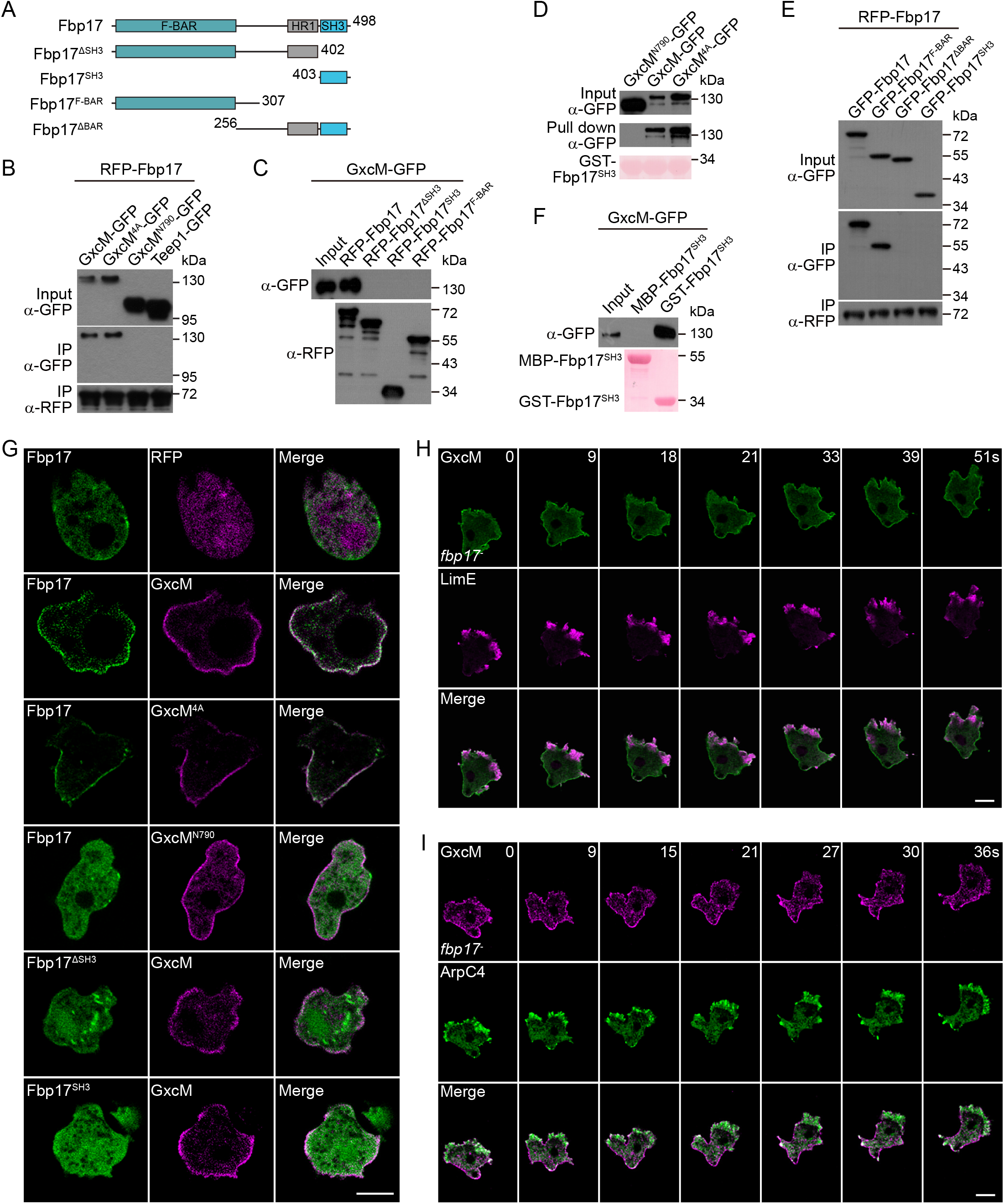
The actin assembly-promoting activity of GxcM relies on interaction with Fbp17. (A) Schematic representation of full-length Fbp17 and truncation constructs. (B) GxcM-GFP and GxcM^4A^-GFP, but not GxcM^N790^-GFP or Teep1-GFP, co-immunoprecipitated with RFP-Fbp17. (C) RFP-Fbp17, but not RFP-Fbp17 truncations, co-immunoprecipitated with GxcM-GFP. (D) GST-Fbp17^SH3^ pulled down GxcM-GFP and GxcM^4A^-GFP, but not GxcM^N790^-GFP. The protein-transferred membrane was stained with Ponceau S to show purified GST fusion proteins. (E) GFP-Fbp17 and GFP-Fbp17^F-BAR^, but not GFP-Fbp17^ΔBAR^ or GFP-Fbp17^SH3^, co-immunoprecipitated with RFP-Fbp17. (F) GST-Fbp17^SH3^, but not MBP-Fbp17^SH3^, pulled down GxcM-GFP. The protein-transferred membrane was stained with Ponceau S to show purified GST and MBP fusion proteins. (G) Localization of GFP-Fbp17 or -Fbp17 truncations and the RFP control or RFP-tagged GxcM, GxcM^4A^, or GxcM^N790^ in randomly migrating cells. (H) Time-lapse imaging of GxcM-GFP and RFP-LimEΔcoil in *fbp17*^−^ cells migrating under agarose along a folic acid gradient. (I) Time-lapse imaging of GxcM-RFP and GFP-ArpC4 in *fbp17*^−^ cells migrating under agarose along a folic acid gradient. Scale bars, 5 μm.

Co-immunoprecipitation experiments were performed to verify the interaction between GxcM and Fbp17. GxcM-GFP, but not Teep1-GFP, co-immunoprecipitated with RFP-Fbp17 (Fig. 3B). GEF domain mutation (GxcM^4A^-GFP) did not affect the interaction, but deletion of the C-terminal region (GxcM^N790^-GFP) abolished the interaction (Fig. 3B), which was consistent with the MS experiment (Fig. S6B). Next, we examined whether the SH3 domain of Fbp17 mediates this interaction. Truncated Fbp17 lacking the SH3 domain (RFP-Fbp17^ΔSH3^ or RFP-Fbp17^F-BAR^) lost the ability to interact with GxcM-GFP (Fig. 3C). Furthermore, purified recombinant glutathione-S-transferase (GST)-Fbp17^SH3^ was able to effectively pull down GxcM-GFP and GxcM^4A^-GFP, but not GxcM^N790^-GFP, from cell lysates (Fig. 3D). However, we noted that RFP-Fbp17^SH3^ could not precipitate GxcM-GFP (Fig. 3C). Because F-BAR proteins tend to form homodimers (Frost et al., 2008; Shimada et al., 2007), we speculated that the ability to dimerize via either the F-BAR domain or GST may be required in binding with GxcM. In support of this notion, we found that Fbp17 likely self-interacts via the F-BAR region (Fig. 3E). Furthermore, unlike GST-Fbp17^SH3^, recombinant maltose-binding protein (MBP)-fused Fbp17^SH3^ failed to pull down GxcM (Fig. 3F). These experiments indicate that GxcM and Fbp17 interact via association of the C-terminus of GxcM with the SH3 domain of Fbp17.

When expressed as a GFP-fusion protein in cells, a small but discernable fraction of GFP-Fbp17 was found at the periphery of cells (Fig. S7, D and E). Deletion of *gxcM* did not evidently change this pattern, but expression of GxcM-RFP or GxcM^4A^-RFP strongly recruited GFP-Fbp17 to the cortex (Fig. 3G). Consistent with the interaction data, expression of GxcM^N790^-RFP could not recruit GFP-Fbp17, nor could GxcM-RFP GFP-Fbp17^ΔSH3^ (Fig. 3G).

To examine the functional significance of the GxcM-Fbp17 interaction, we generated *fbp17* knockout (*fbp17*^−^) cells (Fig. S8A) and examined the consequence of GxcM overexpression in the absence of *fbp17*. Deletion of *fbp17* did not affect the localization of GxcM (Fig. S8, C and D), but it disrupted the actin assembly-promoting activity of GxcM. In contrast to what was observed in GxcM-GFP/WT cells, the ectopic accumulation of LimEΔcoil and AprC4 in the rear of migrating cells (Fig. 1, J and L; Fig. 2, B and D) was nearly completely abolished in GxcM-GFP/*fbp17*^−^ cells (Fig. S8C; Fig. 3, H and I; Video 5), with the bulk of LimEΔcoil and AprC4 signals distributing to the migrating front as in WT cells. These experiments indicate that the actin assembly-promoting activity of GxcM requires association with Fbp17, and Fbp17 likely functions downstream of GxcM to regulate cortical actin generation.

### Fbp17 is required to maintain cortical integrity

We examined whether development, cytokinesis, or migration is affected by *fbp17* deletion. On bacterial lawns, *fbp17*^−^ cells formed smaller plaques but were able to advance through development (Fig. S8B). This phenotype could be fully rescued by expression of GFP-Fbp17 (Fig. S8B). When assayed for cytokinesis, the vast majority of WT and rescue cells were mono- or bi-nucleated. In contrast, *fbp17*^−^ cells exhibited increased failure of cytokinesis; merely 33.5% of cells were mono-nucleated, whereas more than 25% developed three or more nuclei (Fig. 4, A and B). In random motility assays, *fbp17*^−^ cells presented a decrease of approximately 40% in the speed of movement and 70% reduction in the Euclidean distance, indicating that the knockout cells migrated more slowly and less persistently (Fig. 4, C-E). When assayed for under agarose chemotaxis, the *fbp17*^−^ cells migrated towards the folic acid source at a speed comparable to that of WT cells (Fig. 4, F-H). Despite the recovery of speed, the *fbp17*^−^ cells still exhibited reduced persistence and directionality, which was reflected in the more erratic cell tracks and reduced net movement up the gradient (Fig. 4, F-H).

**Figure 4.**
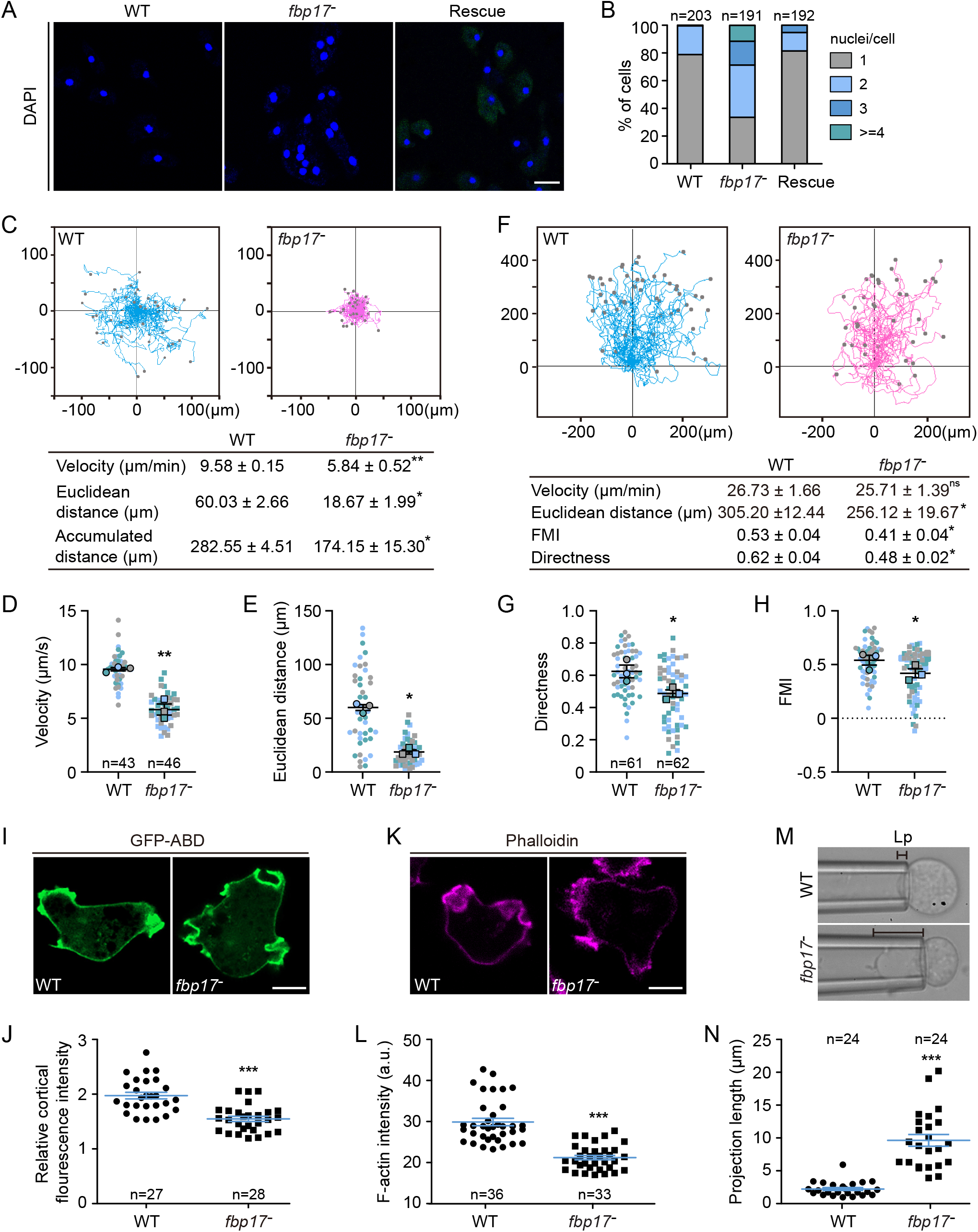
Fbp17 is required to maintain cortical integrity. (A) WT, *fbp17*^−^, and GFP-Fbp17/*fbp17*^−^ (rescue) cells grown for 60 h in shaken suspension were fixed and stained with DAPI to visualize the nuclei. (B) Quantification of nuclei in cells. n, number of cells analyzed. (C) Top: Trajectories of randomly migrating cells (n = 43 for WT and 46 for *fbp17*^−^). Bottom: Summary of motility parameters. (D-E) Velocity and Euclidean distance of cells shown in C. At least 13 cells were quantified per experiment (each experiment shown in a different color); mean ± s.e.m. (F) Top: Trajectories of cells migrating under 2% agarose along a folic acid gradient (n = 61 for WT and 62 for *fbp17*^−^). Bottom: Summary of chemotaxis parameters; FMI, forward migration index. (G-H) Directness and FMI of cells shown in F. At least 17 cells were quantified per experiment (each experiment shown in a different color); mean ± s.e.m. (I) Localization of GFP-ABD in WT and *fbp17*^−^ cells. (J) Quantification of the cortex-to-cytoplasm fluorescent intensity ratios of GFP-ABD. (K) WT and *fbp17*^−^ cells were fixed and stained with Alexa Fluor 555-labeled phalloidin. (L) Quantification of the fluorescence intensity of cortical phalloidin. (M) Projection length (Lp) of WT and *fbp17*^−^ cells determined by micropipette aspiration using a constant pressure of 500 Pa for 5 min. (N) Quantitative analysis of the projection lengths of probed cells. For (C) and (F), data were from three independent experiments; mean ± s.e.m. (the average of each biological replicate was used to calculate the mean and s.e.m.). For (J), (L), and (N), data were from three independent experiments; the scatter plot shows data points with mean ± s.e.m.; n, number of cells analyzed. Scale bars, 5 μm.

To examine whether these morphogenesis and migration defects were caused by defects in cortex assembly, we visualized cortical actin using the ABD120-GFP reporter and phalloidin staining (Pang et al., 1998; Zatulovskiy et al., 2014). In WT cells, the ABD120-GFP signal was more concentrated at the leading-edge protrusions, yet a significant fraction marked the cortical region at the rear and lateral sides (Fig. 4I). Notably, deletion of *fbp17* did not seem to affect the ABD120 signal at the leading edge, but reduced it at the cortical region (Fig. 4I). Quantification revealed a reduction of approximately 20% in the cortical-to-cytosol fluorescent intensity ratio of ABD120-GFP in *fbp17*^−^ cells compared to WT cells (Fig. 4J). Similar results were obtained with phalloidin staining, in which *fbp17*^−^ cells exhibited a decrease of approximately 30% in the cortical intensity of phalloidin (Fig. 4, K and L).

As the F-actin shell is the main contributor to cortical rigidness, we examined whether the apparent reduction in cortical F-actin content in *fbp17*^−^ cells led to a weakened cortex. To this end, we performed micropipette aspiration assays (MPAs) to measure the global mechanical rigidity of suspended cells. We quantified the initial projection length (Lp) of cells captured at a constant pressure of 500 Pa. Under this condition, the plasma membrane deformed along the micropipette, which negatively correlated with the cell’s cortical rigidness (Hochmuth, 2000; Ramalingam et al., 2015). Deletion of *fbp17* resulted in significantly longer projection lengths (mean ± s.e.m.: *fbp17*^−^, 9.64 ± 0.89 μm; WT, 2.23 ± 0.22 µm), implying reduced cortical rigidity (Fig. 4, M and N). A small fraction of *fbp17*^−^ cells were aspirated into the micropipette within 5 min, further validating that the ability to resist mechanical forces was impaired. These results substantiate the role of Fbp17 in cortical actin assembly.

The actin cortex has been shown modulate the subcellular localization of myosin II, a main driver of cell contractility (Vicente-Manzanares et al., 2009). We compared the distribution of GFP-myosin II in WT and *fbp17*^−^ cells. Deletion of *fbp17* reduced the extent of myosin II accumulation in the rear cortical region of randomly migrating cells (Fig. S8E). Placing cells under compression was shown to recruit actin and myosin II to the cortex, which counteracts the applied force (Laevsky and Knecht, 2003; Ramalingam et al., 2015). Consistently, we observed increased accumulation of myosin II in the rear of WT cells chemotaxing under 2% agarose (Fig. S8F). Intriguingly, this condition restored the posterior signal of myosin II in *fbp17*^−^ cells (Fig. S8F). Taking into account the motility measurements acquired under the same conditions (Fig. 4, C-H), these data suggest that a dysfunction in the actomyosin meshwork resulting from *fbp17* deletion may underlie the decreased random motility, whereas adaptation to mechanical stimulation could partially rescue the defect and restore migration.

### Fbp17 promotes WASP-mediated actin polymerization

Given the role of Fbp17 in supporting cortical integrity and mediating the downstream effect of GxcM, as well as the roles of its homologues in other organisms, we speculated that Fbp17 may contribute to Arp2/3-dependent actin polymerization by interacting with and stimulating the activities of WASP family NPFs (Ho et al., 2004; Takano et al., 2008; Tsujita et al., 2006). We performed co-immunoprecipitation experiments to examine whether Fbp17 interacts with WASP, which is encoded by a single gene, *wasA*, in *Dictyostelium*. GFP-WASP, but not Teep1-GFP, co-immunoprecipitated with RFP-Fbp17 (Fig. 5A). The interaction depended on the SH3 domain of Fbp17, as RFP-Fbp17^ΔSH3^ lost the ability to interact with WASP (Fig. 5B). Interestingly, as observed for the Fbp17-GxcM interaction (Fig. 3, C and D), the interaction between Fbp17 and WASP also appeared to require the presence of a potentially dimerized SH3. Fbp17^SH3^ expressed in cells or as MBP-fusion protein could not precipitate GFP-WASP, whereas GST-Fbp17^SH3^ pulled down GFP-WASP efficiently (Fig. 5C). Furthermore, consistent with the proposed links among GxcM, Fbp17, and WASP, a fraction of GFP-WASP was recruited to cortical regions with intensive GxcM-RFP signals, and this recruitment was abolished by *fbp17* deletion (Fig. 5D).

**Figure 5.**
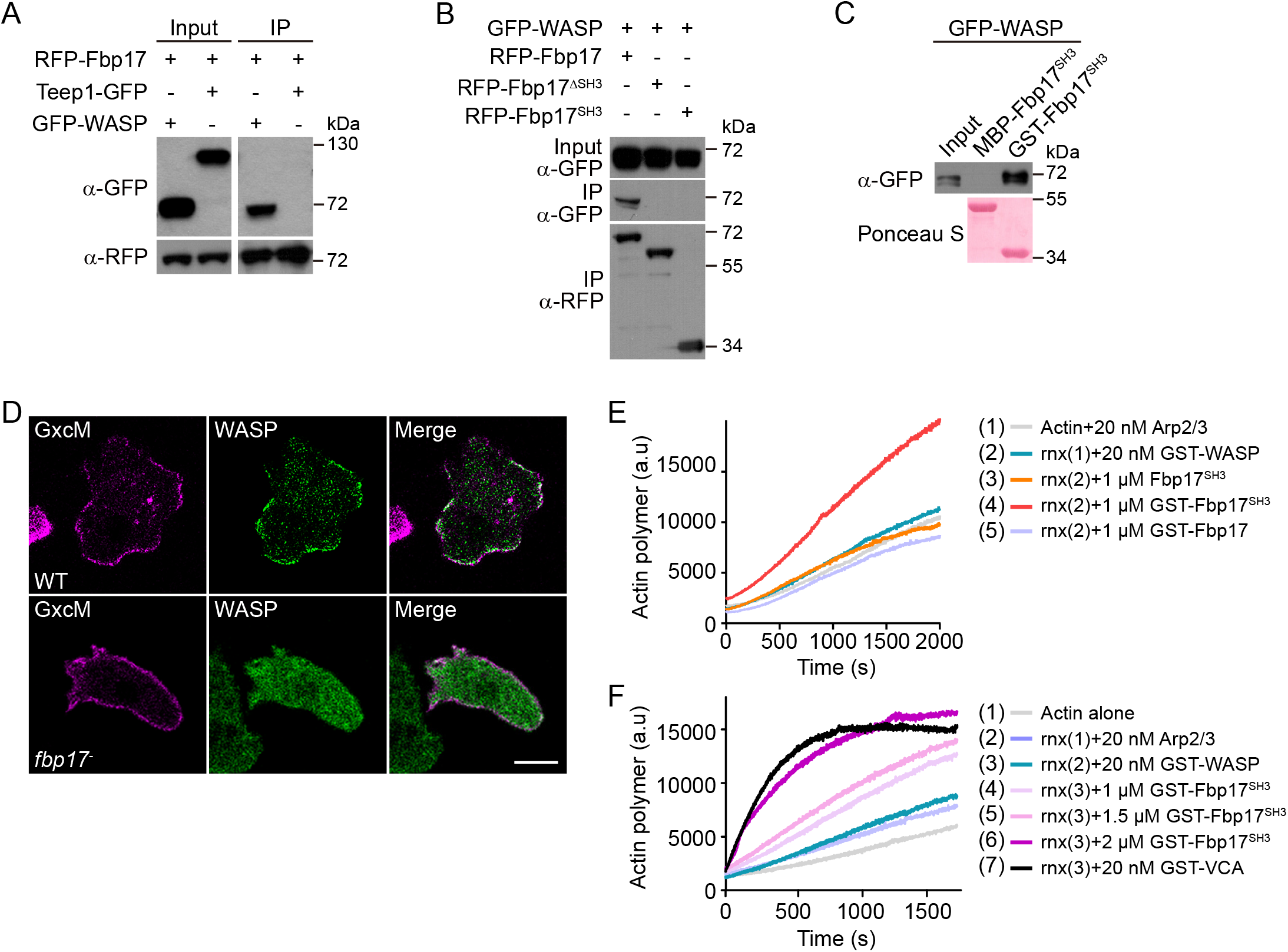
Fbp17 promotes WASP-mediated actin polymerization. (A) GFP-WASP, but not Teep1-GFP, co-immunoprecipitated with RFP-Fbp17. (B) RFP-Fbp17, but not -Fbp17^ΔSH3^ or - Fbp17^SH3^, co-immunoprecipitated with GFP-WASP. (C) GST-Fbp17^SH3^, but not MBP-Fbp17^SH3^, pulled down GFP-WASP. The protein-transferred membrane was stained with Ponceau S to show purified GST and MBP fusion proteins. (D) Expression of GxcM-RFP in WT but not *fbp17*^−^ cells triggered the cortical recruitment of GFP-WASP. Scale bar, 5 μm. (E) GST-Fbp17^SH3^, but not Fbp17^SH3^ or GST-Fbp17, promotes WASP- and Arp2/3-mediated actin polymerization in pyrene assays. (F) GST-Fbp17^SH3^ promotes WASP- and Arp2/3-dependent actin polymerization in a concentration-dependent manner. The VCA domain of WASP purified as a GST fusion was included as a control.

By the in vitro pyrene-actin polymerization assay, we measured the activity of recombinant Fbp17 toward WASP. When mixing with actin and the Arp2/3 complex, GST-WASP purified from insect cells exhibited minimal actin polymerization-promoting activity, indicating that it likely adopted an autoinhibitory configuration (Fig. 5E). Adding purified Fbp17^SH3^ or GST-Fbp17 did not significantly change the kinetics of the reaction (Fig. 5E). In contrast, purified GST-Fbp17^SH3^ promoted actin polymerization driven by WASP and Arp2/3 in a concentration-dependent manner (Fig. 5, E and F). The VCA domain of *Dictyostelium* WASP (319-399 aa) was included as a positive control (Fig. 5F). Taken together, these experiments indicate that Fbp17 likely facilitates WASP activation downstream of GxcM in cortical actin formation.

### RacC acts downstream of GxcM to regulate cortex assembly via the activation of WASP

Our results described thus far suggest a scenario in which the rear cortex-localized GxcM recruits Fbp17 to promote WASP and Arp2/3-mediated actin assembly. However, as presented earlier, the actin assembly-promoting activity of GxcM also requires its Rho GEF domain, implying involvement of Rho GTPase(s) in this process. *Dictyostelium* cells lack canonical Cdc42 and Rho homologs but express 20 Rac proteins, some of which exhibit characteristics of Cdc42 and Rho (Filic et al., 2021). We investigated whether one or more of these Rac proteins act downstream of GxcM. Our MS analyses of GxcM-associated proteins (Fig. S6B) did not yield a possible candidate. Considering that Rho GTPases are potent regulators of WASP and the proposed mechanism by which FBP17 family proteins function in other organisms involves association with Rho GTPases (Ho et al., 2004; Watson et al., 2016; Watson et al., 2017), we decided to seek the relevant small GTPases by looking for binding partners of WASP and Fbp17.

We employed yeast two-hybrid (Y2H) analyses to screen Rac proteins for interaction with the GTPase-binding domain (GBD, aa 126-230) of WASP or full-length Fbp17 (Fig. 6A; Fig. S9A). The GBD domain interacted with several Racs (Rac1A, Rac1B, Rac1C, RacA, RacB, and RacC) in their constitutively active (CA) forms, whereas Fbp17 interacted selectively with the CA form of RacC (RacC^CA^). RacE, which has been shown to regulate cortex assembly via formin proteins (Litschko et al., 2019), did not exhibit interaction with either GBD or Fbp17 (Fig. S9A). We focused our subsequent investigations on RacC because of three reasons: (1) In line with the notion that GEFs preferentially bind to the nucleotide-free form of small GTPases, we found that RacC, but not Rac1A, exhibited an EDTA-dependent interaction with GxcM (Fig. 6B); (2) a previous study using a cell-free system showed that GTPγS-charged RacC is capable of stimulating F-actin polymerization via the activation of WASP (Han et al., 2006); (3) overexpression of RacC has been shown to induce unusual actin-based structures, which we found somewhat resemble the effect of GxcM overexpression (Seastone et al., 1998).

**Figure 6.**
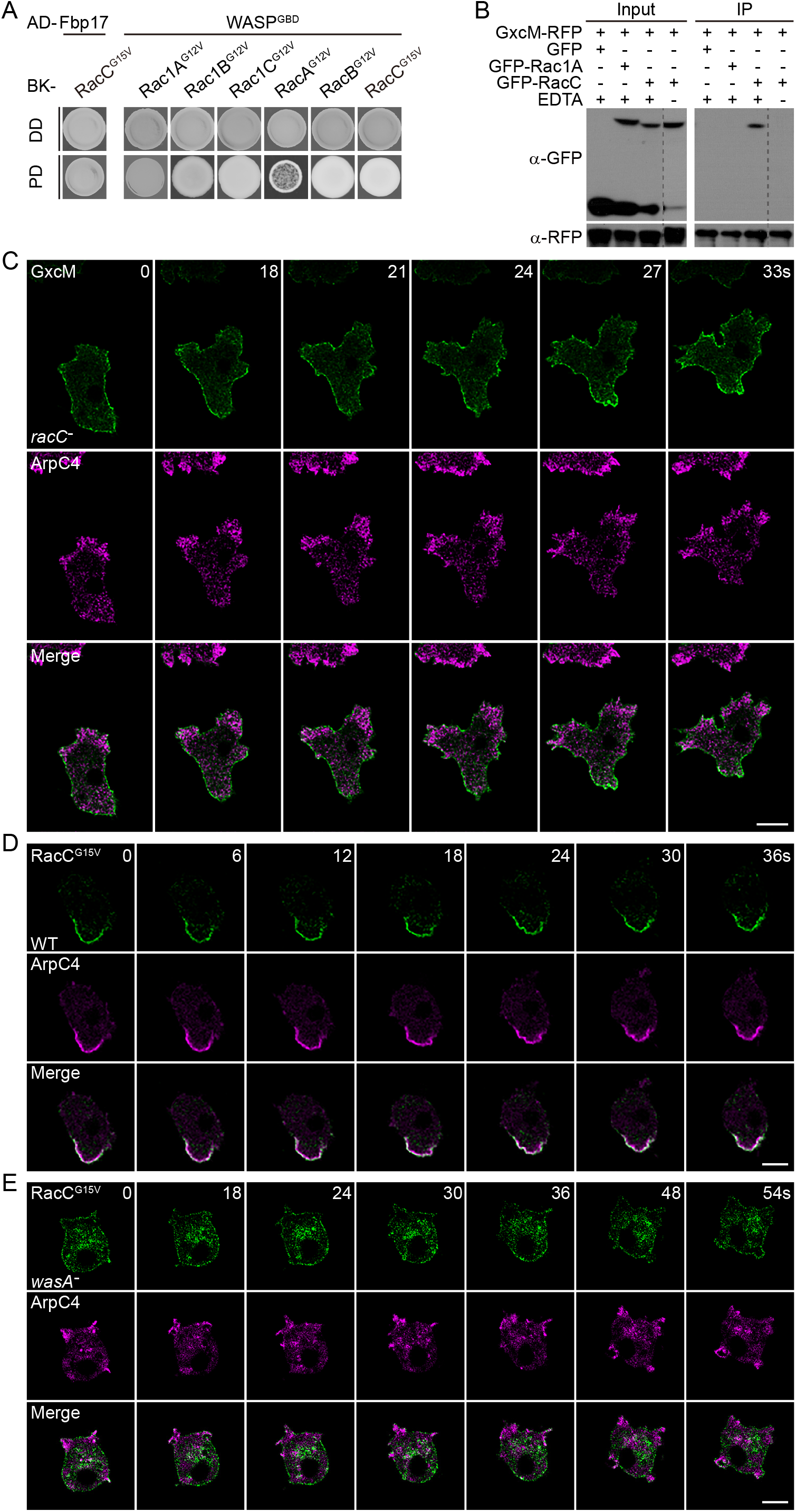
RacC acts downstream of GxcM to regulate cortical actin assembly via the activation of WASP. (A) Yeast two-hybrid assay showing that the interaction between Fbp17 or the GBD domain of WASP with the constitutively active (CA) forms of the indicated Rac proteins. (B) GFP-RacC, but no GFP-Rac1A, co-immunoprecipitated with GxcM-RFP in the presence of EDTA. (C) Time-lapse imaging of GxcM-GFP and RFP-ArpC4 in *racC*^−^ cells migrating under agarose along a folic acid gradient. (D) Time-lapse imaging of GFP-RacC^G15V^ and RFP-ArpC4 in randomly migrating WT cells. (E) Time-lapse imaging of GFP-RacC^G15V^ and RFP-ArpC4 in randomly migrating *wasA*^−^ cells. Scale bars, 5 μm.

By generating cells lacking *racC* (Fig. S9B) or expressing the CA forms of RacC (RacC^G15V^ or RacC^Q64L^), we further investigated the role of RacC in cortex assembly in relation to GxcM, Fbp17, and WASP. As with the deletion of *fbp17*, deletion of *racC* effectively blocked GxcM overexpression-induced rear accumulation of AprC4 or LimEΔcoil in directional migrating cells without disrupting the localization of GxcM (Fig. 6C; Fig. S9D; Video 6). This result indicates that RacC is indeed necessary for GxcM-mediated actin polymerization. Intriguingly, expression of GFP-RacC^G15V^ under an inducible promoter generated increasingly polarized cells with the RacC^G15V^ signal selectively localized to the rear side (Fig. 6D; Fig. S10A; Video 7). In these cells, ArpC4 and LimEΔcoil redistributed from the migrating front to where GFP-RacC^G15V^ was most concentrated. Expression of GFP-RacC^Q64L^ caused a similar redistribution of the actin markers (Fig. S10, B and C). Therefore, although the expression of RacC^CA^ did not fully recapitulate the effect of GxcM overexpression, it transformed the cortical actin organization in a similar manner. Consistent with the Y2H result, the expression of RFP-RacC^CA^ was able to recruit both GFP-WASP and GFP-Fbp17 to the cell cortex (Fig. S10D). Furthermore, RacC^CA^-induced polarization and actin reorganization were abolished by *wasA* deletion, but not *fbp17* deletion, placing RacC upstream of WASP in the network (Fig. 6E; Fig. S10E; Fig. S11A). Taken together, these results indicate that, similar to Fbp17, RacC functions as an intermediate connecting GxcM and WASP activities to regulate cortical actin assembly.

We examined whether deletion of *racC* impairs cortical function. The *racC*– cells formed smaller plaques on bacterial lawns (Fig. S9B), exhibited a severe defect in cytokinesis (Fig. 7, A and B), migrated with reduced speed and directionality (Fig. 7, C-I), and failed to properly localize myosin II (Fig. S9E), indicating that cortex-dependent cellular activities were all affected to varying degrees by *racC* deletion. The cortical F-actin content and global mechanical rigidity of cells measured in the MPA (projection length mean ± s.e.m.: *racC*^-^, 8.70 ± 0.76 μm; WT, 2.25 ± 0.15 µm) were also clearly impaired in *racC*^−^ cells compared to control (Fig. 7, J-O), further substantiating the role of RacC in cortical activity regulation.

**Figure 7.**
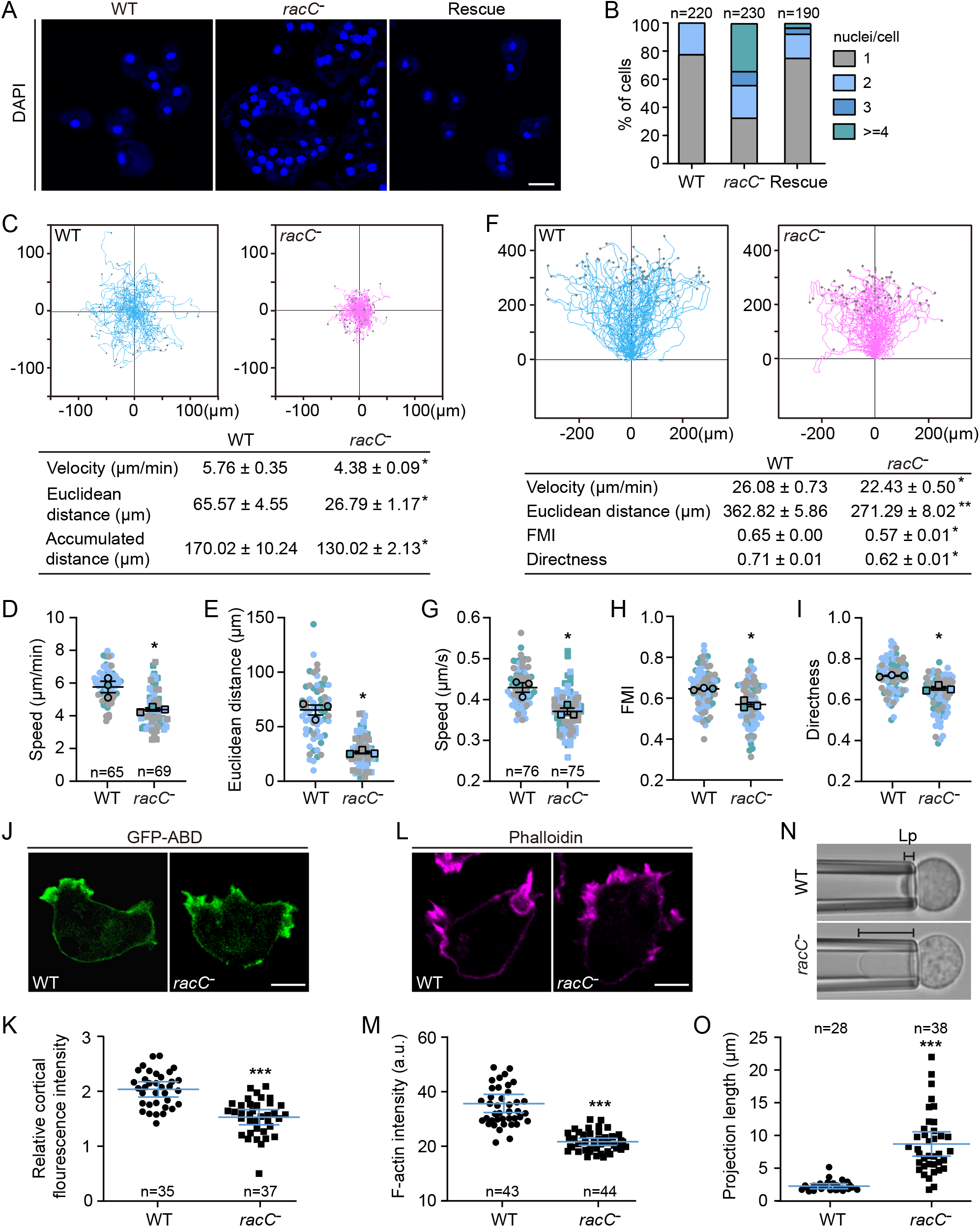
RacC is required to maintain cortical integrity. (A) WT, *racC*^−^, and GFP-RacC/*racC*^−^ (rescue) cells grown for 60 h in shaken suspension were fixed and stained with DAPI to visualize the nuclei. (B) Quantification of nuclei in cells. n, number of cells analyzed. (C) Top: Trajectories of randomly migrating cells (n = 65 for WT and 69 for *racC*^−^). Bottom: Summary of motility parameters. (D-E) Velocity and Euclidean distance of cells shown in C. At least 15 cells were quantified per experiment (each experiment shown in a different color); mean ± s.e.m. (F) Top: Trajectories of cells migrating under 2% agarose along folic acid gradient (n = 76 for WT and 75 for *racC*^−^). Bottom: Summary of chemotaxis parameters. (G-I) Speed, FMI, and Directness of cells shown in F. At least 20 cells were quantified per experiment (each experiment shown in a different color); mean ± s.e.m. (J) Localization of GFP-ABD in WT and *racC*^−^ cells. (K) Quantification of the cortex-to-cytoplasm fluorescent intensity ratios of GFP-ABD. (L) WT and *racC*^−^ cells were fixed and stained with Alexa Fluor 555-labeled phalloidin. (M) Quantification of the fluorescence intensity of cortical phalloidin. (N) Projection length (Lp) of WT and *racC*^−^ cells determined by micropipette aspiration using a constant pressure of 500 Pa for 5 min. (O) Quantitative analysis of the projection lengths of probed cells. For (C) and (F), data were from three independent experiments; mean ± s.e.m. (the average of each biological replicate was used to calculate the mean and s.e.m.). For (K), (M), and (O), data were from three independent experiments; the scatter plot shows data points with mean ± s.e.m.; n, number of cells analyzed. Scale bars, 5 μm.

Finally, we found that deletion of *wasA* suppressed the effect of GxcM overexpression. Although cell morphology and polarity were not fully restored as seen in *fbp17*^−^ or *racC*– cells overexpressing GxcM, the intense GxcM-induced signals of peripheral actin patches were no longer present in *wasA*^−^ cells overexpressing GxcM (Fig. S11C). In addition, the *wasA*– cells exhibited considerably reduced cortical rigidity measured in the MPA (projection length mean ± s.e.m.: *wasA*^−^, 8.39 ± 0.71 μm; WT, 1.96 ± 0.20 µm; Fig. S11D). Deletion of *wasA* has been associated with defects in multicellular development, cytokinesis, myosin recruitment, and migration (Davidson et al., 2018), which is in line with a critical role in cortex function proposed in our study. Taken together, our results delineate a novel signaling pathway contributing to the generation of the rear cortical actin subcompartment in migrating *Dictyostelium* cells.

## Discussion

The cell cortex has been shown to be composed of both formin-mediated linear F-actin and Arp2/3-mediated branched F-actin. However, most previous studies have used relatively non-motile cells, such as mammalian cells constitutively blebbing, rounded up for mitosis, or detached from the substratum, as model systems (Bovellan et al., 2014; Cao et al., 2020; Chugh et al., 2017). Here, by analyzing cortical actin assembly in *Dictyostelium* cells, we show that a signaling cascade promotes Arp2/3-dependent actin polymerization and reinforces cortical integrity in the rear of polarized migrating cells. In this cascade, GxcM localized selectively to the rear of cells signals through Fbp17 and RacC, which in turn activate WASP to stimulate actin polymerization (Fig. S12). Together with reports showing the critical role of formin proteins in cortex assembly in *Dictyostelium* (Litschko et al., 2019), these findings indicate that the actin meshwork in the rear cortex of rapidly migrating cells may also be composed of structurally and biochemically different actin filament arrays.

Prior to our study, sporadic observations have hinted at the presence of Arp2/3 and branched F-actin in the rear of *Dictyostelium* cells. However, because the Arp2/3 complex is a main driver of actin-based structures, including pseudopods and macropinocytic cups, live cell reporters for Arp2/3 activity and F-actin are generally highly enriched in the front of cells, likely overshadowing the potential signals in the rear. We were able to visualize and uncover a previously unrecognized function of Arp2/3-based actin in the rear of the cell by overexpressing GxcM or RacC^CA^. Although these were considered non-physiological conditions, the fact that deletion of the respective downstream components of the pathway abolished or reduced the effect of overexpressing GxcM or RacC^CA^ corroborated the functional involvement of endogenous Fbp17, RacC, and WASP. Defects in the various cortex-dependent cellular activities observed in *fbp17*^−^, *racC*^−^, and *wasA*^−^ cells (Davidson et al., 2018) further substantiated the function of these proteins in organizing the cortical actin network. Thus, Arp2/3-mediated F-actin likely not only contributes to the structure of the cortex, but also its function.

A key function of the rear cortex is to mediate retraction. The myosin II contractile machinery often uses relatively long actin filaments assembled by the formin family proteins (Svitkina, 2020). A previous study in *Dictyostelium* showed that formin mutants failed to properly polarize and recruit myosin II (Litschko et al., 2019). Our data suggest that the Arp2/3-mediated branched actin network may also contribute to the subcellular localization of myosin II. Cells deleted of *fbp17* failed to robustly recruit GFP-myosin II to the rear, though compression of the cells under agarose, a condition known to strengthen the cortex by relocating actin, myosin II, and other actin-binding proteins, rescued the defect. Similar defects in myosin II localization were observed in *racC*^−^ and *wasA*^−^ cells (Davidson et al., 2018). Future studies are needed to elucidate how different actin filament arrays, together with other signaling and mechanical cues (Bosgraaf and van Haastert, 2006), fine-tune the activity and localization of myosin II.

Why generation of the cortex requires the combined action of different actin filaments and, thus, different nucleators represents an intriguing question. Studies in HeLa and M2 cells have shown that perturbation of the formin mDia1 and Arp2/3 complex leads to decreases in cortical F-actin but distinct impacts on the nanoscale architecture and mechanical properties of the network (Bovellan et al., 2014). Interestingly, the RacE-ForA/ForE/ForH pathway and the GxcM-Fbp17/RacC-WASP pathway also appear to contribute differently to the cortical actin network. The triple formin and *racE* mutants contained large peripheral patches devoid of F-actin and exhibited severe defects in cortical rigidity, cytokinesis, and development. They moved with exaggerated fronts and higher speed in unconfined environments but nearly completely lost the ability to migrate in 2D-confinement (Litschko et al., 2019). In contrast, deletion of *fbp17* or *racC* caused relatively milder defects in cortical actin content and rigidity, cytokinesis, and development. The mutant cells were able to migrate in 2D-confinement, but moved with significantly slower speed in unconfined settings. These observations highlight the functional difference between the formin-mediated and Arp2/3-mediated actin structures. Different nucleators likely differentially affect the length, degree of branching, and density of cortical actin filaments, all of which influence the emerging physical properties of the cortical network and, thus, cell behavior (Chugh et al., 2017; Ennomani et al., 2016; Fritzsche et al., 2016; Koenderink and Paluch, 2018). Synergistic action among distinct nucleators has also been observed (Cao et al., 2020). It will be of great interest in future studies to understand the relative contribution of different actin nucleator to the rear cortex and their interplay in cortical function.

Our analysis of the GxcM-Fbp17/RacC-WASP signaling cascade revealed complex crosstalk and redundancy. First, GxcM appears to promote WASP activity by recruiting Fbp17, as well as activating RacC. Both branches of the cascade are necessary for the full activity, as disruption of either blocked the effect of GxcM overexpression. However, the two branches are not independent, but likely coordinate with each other. Fbp17 interacts with RacC in its GTP-bound form, likely through the Rho-binding HR1 domain. As the HR1 domain in the FBP17 family of proteins generally exhibits a much lower affinity for Rho GTPase compared to the WASP GBD domain, an effector handover model has been proposed (Watson et al., 2016; Watson et al., 2017). We speculate that Fbp17 and RacC work in a similar manner to activate WASP (Fig. S12): GxcM recruits Fbp17 to where RacC is activated; homodimerization, and possibly oligomerization, of Fbp17 (Frost et al., 2008; Shimada et al., 2007; Snider et al., 2021) results in a sufficient local concentration to achieve interaction with activated RacC; Fbp17 further recruits WASP, and the positioning of WASP by Fbp17, along with a potential allosteric effect, allows the GBD domain to contact RacC; and the high affinity of GBD for RacC ensures that an unfolded state of WASP is favored, promoting the release of VCA, which ultimately triggers robust Arp2/3 activation and actin nucleation. Second, although overexpression of GxcM strongly activated the signaling cascade, its deletion caused only subtle defects (Fig. S2), suggesting that additional factors, possibly other RhoGEF proteins, may substitute for GxcM in cells. Third, unlike deletion of *fbp17* or *racC*, deletion of *wasA* only partially suppressed the effect of GxcM overexpression, implying redundancy in the output of the cascade. In addition to WASP, FBP17 family proteins have been shown to act through the SCAR/WAVE complex (Bai and Grant, 2015). Notably, a previous study using the *Dictyostelium* SCAR complex as bait identified Fbp17 as an interacting partner (Fort et al., 2018). Therefore, future studies are needed to investigate the system redundancy and elucidate the precise biochemical mechanism by which the cortical branched actin network is assembled.

Another open question is how the GxcM-Fbp17/RacC-WASP signaling cascade is selectively brought to the rear of the cell. GxcM, which is positioned in the upstream of the pathway, likely plays an essential role. GxcM exhibits dynamic behavior identical to other rear proteins, including dissociating from the protrusions and responding to global chemoattractant stimulation by transiently falling off the membrane (Iijima and Devreotes, 2002; Swaney et al., 2015). We previously showed that back-to-front gradients of PI(4,5)P_2_ and PI(3,5)P_2_ jointly regulate the rear accumulation of a number of proteins, such as Teep1 (Li et al., 2022). However, GxcM did not exhibit apparent PI(4,5)P_2_- or PI(3,5)P_2_-binding activity assessed in the lipid dot blot assay. Thus, the molecular mechanism underlying its rear accumulation remains obscure. Intriguingly, activated RacC also selectively targets to the rear of the cell, similar to activated RacE as reported previously (Litschko et al., 2019; Senoo et al., 2020). We found that, for RacC, such selectivity depends on the downstream activity of the pathway and is abrogated by removing *wasA* from the system. Thus, GxcM-mediated specific targeting and feedback from actin polymerization may act together to maintain the polarized distribution of the signaling cascade.

## Figure Legend

**Figure S1. GxcM exhibits characteristics of a rear protein**. (A) Translocation of GxcM-GFP in response to folic acid stimulation added at time 0 in the presence of 5 μM LatA (pre: before stimulation). (B) Translocation of GxcM^N790^-GFP in response to cAMP stimulation in the presence of LatA. (C-D) Localization of GxcM^N455^-GFP and GxcM^N456-770^-GFP in randomly migrating cells. (E) Translocation of GxcM^4A^-GFP in response to cAMP stimulation in the presence of LatA. (F) Sequence alignment of the DH domains of GxcM and human ECT2. The red box indicates the conserved residues shown to be essential for the GEF activity of ECT2. Scale bars, 5 μm.

**Figure S2. Characterization of *gxcM* deletion cells**. (A) Top: Design of the knockout construct. A blasticidin resistant cassette (BSR) was inserted to replace part of the open reading frame of *gxcM*. Bottom: Targeted clones were confirmed by PCR using the indicated primers. (B) WT and *gxcM*^−^ cells were plated clonally with bacteria (*Klebsiella aerogenes*) on standard medium agar for 5 d. Scale bar, 5 mm. (C) Top: WT and *gxcM*^−^ cells grown for 60 h in shaken suspension were fixed and stained with DAPI to visualize the nuclei. Bottom: Quantification of nuclei in cells. n, number of cells analyzed. Scale bar, 5 μm. (D) Top: Trajectories of randomly migrating cells (n = 38 for WT and 47 for *gxcM*^−^). Bottom: Summary of motility parameters. Data were from three independent experiments; mean ± s.e.m. (the average of each biological replicate was used to calculate the mean and s.e.m.). (E) Top: Trajectories of cells migrating under 2% agarose along a folic acid gradient (n = 41 for WT and 41 for *gxcM*^−^). Bottom: Summary of chemotaxis parameters. Data were from three independent experiments; mean ± s.e.m. (the average of each biological replicate was used to calculate the mean and s.e.m.). (F) Left: Projection length (Lp) of WT and *gxcM*^−^ cells determined by micropipette aspiration using a constant pressure of 500 Pa for 5 min. Right: Quantitative analysis of the projection lengths of probed cells. Data were from one representative experiment; the scatter plot shows data points with mean ± s.e.m.; n, number of cells analyzed.

**Figure S3. Characterization of cells overexpressing GxcM**. (A) Time-lapse imaging of cells expressing PHcrac-GFP and Teep1-RFP. (B) Time-lapse imaging of cells expressing PHcrac-GFP and GxcM-RFP. (C) Top: Trajectories of randomly migrating cells (n = 64 for GFP/WT and 50 for GxcM-GFP/WT). Bottom: Summary of motility parameters. Data were from three independent experiments; mean ± s.e.m. (D) Quantification of TRITC dextran (TD) uptake. Data were from three independent experiments; the scatter plot shows data points with mean ± s.e.m.; n, number of cells analyzed. (E) Cells expressing GxcM-GFP from an expression cassette integrated into the genome as a stable single copy via restriction enzyme-mediated integration (REMI) was immunostained with an anti-GFP antibody. (F) Time-lapse imaging of randomly migrating cells expressing GFP-ArpC4 and untagged GxcM. (G) Left: GFP/WT and GxcM-GFP/WT cells grown on cell culture plate were fixed and stained with DAPI to visualize the nuclei. Right: Quantification of nuclei. n, number of cells analyzed. Scale bars, 5 μm.

**Figure S4. GxcM overexpression induces actin polymerization and impairs cell motility**. (A-B) Localization of GFP-Coronin and RFP (A) or GxcM-RFP (B) in WT cells migrating under agarose along a folic acid gradient. (C) Top: Trajectories of cells migrating under 0.5% agarose along a folic acid gradient (n = 30 for GFP/WT and 29 for GxcM-GFP/WT). Bottom: Summary of chemotaxis parameters. Data were from at least three movies; means ± s.e.m. Scale bars, 5 μm.

**Figure S5. The cortical actin assembly-promoting activity of GxcM requires the GEF domain as well as an intact C-terminus**. (A, C, E) Localization of GxcM^4A^-RFP with GFP-LimEΔcoil, -Coronin, and -ArpC4 in randomly migrating WT cells. (B, D, F) Localization of GxcM^N790^-RFP with GFP-LimEΔcoil, -Coronin, and -ArpC4 in randomly migrating WT cells. (G) Time-lapse imaging of GxcM^4A^-RFP and GFP-ArpC4 in WT cells migrating under agarose along a folic acid gradient. (H) Time-lapse imaging of GxcM^N790^-RFP and GFP-ArpC4 in WT cells migrating under agarose along a folic acid gradient. Scale bars, 5 μm.

**Figure S6. Identification of Fbp17 as a binding partner of GxcM**. (A) Sequence of the C-terminal region GxcM (aa 791-1145). Proline residues are highlight in red. The blue underlines indicate linear motifs predicted to bind to SH3 domain by the LMDIPred web server. (B-C) Proteomic identification of Fbp17 as a binding partner of GxcM. The bottom table shows the identified unique peptides of Fbp17 from the GxcM-GFP pull-down.

**Figure S7. Fbp17 is a homolog of the FBP17 family of F-BAR proteins**. (A) Sequence alignment of Fbp17 with human CIP4, FBP17, and TOCA-1. Magenta, green, and blue boxes indicate the F-BAR, homology region 1 (HR1), and SH3 domain, respectively. The HR1 domains of all four proteins contain an antiparallel coiled-coil fold. (B) List of protein domains in Fbp17 analyzed using the Conserved Domain Database. (C) Three-dimensional structure of Fbp17 deduced by the online Robetta service (https://robetta.bakerlab.org/). (D) Time-lapse imaging of GFP-Fbp17 in randomly migrating cells. (E) Time-lapse imaging of GFP-Fbp17 in cells moving under agarose along a folic acid gradient. Scale bars, 5 μm.

**Figure S8. Generation and characterization of *fbp17* deletion cells**. (A) Top: Design of the knockout construct. Bottom: Targeted clones were confirmed by PCR. (B) The indicated cells were plated clonally with bacteria (*Klebsiella aerogenes*) on standard medium agar for 5 d. Scale bar, 5 mm. (C) Localization of GxcM-GFP and RFP-LimEΔcoil or GxcM-RFP and GFP-ArpC4 in randomly migrating *fbp17*^−^ cells. (D) Translocation of GxcM-GFP in *fbp17*^−^ cells in response to cAMP stimulation in the presence of LatA. (E) Left: Localization of GFP-myosin II in randomly migrating cells. Right: Quantification of the relative cortical localization of GFP-myosin II. Data were from three independent experiments; the scatter plot shows data points with mean ± s.e.m.; n, number of cells analyzed. (F) Left: Localization of GFP-myosin II in cells migrating under 2% agarose along a folic acid gradient. Right: Quantification of the relative cortical localization of GFP-myosin II. Data were from three independent experiments; the scatter plot shows data points with mean ± s.e.m.; n, number of cells analyzed. Scale bars, 5 μm.

**Figure S9. RacC interacts with Fbp17 and WASP and acts downstream of GxcM**. (A) Yeast two-hybrid assay showing the interaction between Fbp17 and the GBD domain of WASP with the CA forms of 19 Rac proteins. (B) Top: Design of the *racC* knockout construct. Bottom: Targeted clones were confirmed by PCR. (C) The indicated cells were plated clonally with bacteria (*Klebsiella aerogenes*) on standard medium agar for 5 d. Scale bar, 5 mm. (D) Time-lapse imaging of GxcM-GFP and RFP-LimEΔcoil in *racC*^−^ cells migrating under agarose along a folic acid gradient. (E) Left: Localization of GFP-myosin II in randomly migrating WT and *racC*^−^ cells. Right: Quantification of the relative cortical localization of GFP-myosin II. Data were from three independent experiments; the scatter plot shows data points with mean ± s.e.m.; n, number of cells analyzed. Scale bars, 5 μm.

**Figure S10. Expression of constitutively active RacC induces cortical actin polymerization**. (A) Time-lapse imaging of GFP-RacC^G15V^ and RFP-LimEΔcoil in randomly migrating WT cells. (B) Time-lapse imaging of GFP-RacC^Q64L^ and RFP-LimEΔcoil in randomly migrating WT cells. (C) Time-lapse imaging of GFP-RacC^Q64L^ and RFP-ArpC4 in randomly migrating WT cells. (D) Expression of RFP-RacC^G15V^ recruits GFP-Fbp17 and GFP-WASP to the cell cortex. (E) Time-lapse imaging of GFP-RacC^G15V^ and RFP-ArpC4 in randomly migrating *fbp17*^−^ cells. Scale bars, 5 μm.

**Figure S11. Characterization of *wasA* deletion cells**. (A) Top: Design of the knockout construct. Bottom: Targeted clones were confirmed by PCR. (B) The indicated cells were plated clonally with bacteria (*Klebsiella aerogenes*) on standard medium agar for 5 d. Scale bar, 5 mm. (C) Localization of RFP-GxcM and GFP-ArpC4 in *wasA*^−^ cells. Scale bar, 5 μm. (D) Left: Projection length (Lp) of WT and *wasA*^−^ cells determined by micropipette aspiration using a constant pressure of 500 Pa for 5 min. Right: Quantitative analysis of the projection lengths of probed cells. Data were from two independent experiments; the scatter plot shows data points with mean ± s.e.m.; n, number of cells analyzed.

**Figure S12. The GxcM-Fbp17/RacC-WASP signaling cascade in the formation of a rear cortical subcompartment**. GxcM, which localizes selectively in the rear of *Dictyostelium* cells, signals through Fbp17 and RacC to activate WASP by releasing its VCA tail, which in turn promotes Arp2/3 complex-mediated cortical actin assembly. Disruption of the signaling cascade causes defects in cortical rigidity and function, whereas over-activation of the cascade leads to excessive actin polymerization in the rear of the cell.

## Video Caption

**Video 1** – Localization of GxcM-GFP in WT cells migrating under agarose along a folic acid gradient. Corresponds to figure 1H. Images were captured at 2-s intervals and played back at 24 frames per second. Scale bar = 5 μm.

**Video 2** – Localization of PHcrac-GFP and RFP-tagged GxcM, GxcM^N790^, GxcM^4A^, or Teep1 in WT cells during random migration. Corresponds to figures 1B, 1D, 1F, S3A, and S3B. Images were captured at 6-s intervals and played back at 12 frames per second. Scale bar = 5 μm.

**Video 3** – Localization of LimEΔcoil-RFP and GFP-ArpC4 in WT cells expressing GxcM-GFP, GxcM-RFP, or a control fluorescent protein during random migration. Corresponds to figures 1I-1L. Images were captured at 6-s intervals and played back at 12 frames per second. Scale bar = 5 μm.

**Video 4** – Localization of LimEΔcoil-RFP and GFP-ArpC4 in WT cells expressing GxcM-GFP, GxcM-RFP, or a control fluorescent protein migrating under agarose along a folic acid gradient. Corresponds to figures 2A-2D. Images were captured at 3-s intervals and played back at 6 frames per second. Scale bar = 5 μm.

**Video 5** – Localization of GxcM-GFP/LimEΔcoil-RFP or GxcM-RFP/GFP-ArpC4 in *fbp17*^−^ cells migrating under agarose along a folic acid gradient. Corresponds to figures 3H and 3I. Images were captured at 3-s intervals and played back at 6 frames per second. Scale bar = 5 μm.

**Video 6** – Localization of GxcM-GFP with LimEΔcoil-RFP or RFP-ArpC4 in *racC*^−^ cells migrating under agarose along a folic acid gradient. Corresponds to figures 6C and S9D. Images were captured at 3-s intervals and played back at 6 frames per second. Scale bar = 5 μm.

**Video 7** – Localization of GFP-RacC^G15V^ with LimEΔcoil-RFP or RFP-ArpC4 in WT cells during random migration. Corresponds to figures 6D and S10A. Images were captured at 6-s intervals and played back at 12 frames per second. Scale bar = 5 μm.

## Materials and methods

### Cell culture, transformation, and differentiation

WT cells were derived from Ax2 (Ka) axenic strain provided by Robert Kay laboratory or Ax3 axenic strain provided by Peter Devreotes laboratory. The deletion cell line for *wasA* was generated in Ax3 and all the other deletion cell lines were generated in Ax2. WT and gene deletion cells were cultured in HL5 medium (Formedium; cat# HLF3) supplemented with antibiotics at 22°C following routine procedures. Cells carrying expression constructs were transformed by electroporation and maintained in HL5 containing G418 (10-40 μg/ml), Hygromycin (50 μg/ml), or both as needed. To induce the expression of constitutively active RacC 20 μg/ml doxycycline was added and incubated for 16 h. For differentiation, cells grown in HL5 were washed with development buffer (DB; 5 mM Na_2_HPO_4_, 5 mM KH_2_PO_4_, 2 mM MgSO_4_, and 0.2 mM CaCl_2_), starved in DB for 1 h, and pulsed with 100 nM cAMP every 6 min for 3-5 h.

### Gene disruption and plasmid construction

Plasmids and primers used in this study are listed in Table 1. To make knockout constructs for *gxcM, fbp17, racC*, and *wasA* deletion, a blasticidin S resistance (BSR) cassette (Kimmel and Faix, 2006) was inserted into pBlueScript II SK+ to generate pBlueScript-BSR. 5’ and 3’ arms were PCR-amplified from genomic DNA with primers listed in Table 1 and cloned upstream and downstream of the BSR cassette, respectively. The resulting disruption cassette was amplified by PCR and electroporated into Ax2 or Ax3. Gene disruption was confirmed by resistance to blasticidin (10 μg/ml), PCR, and rescue experiment.

To generate constructs expressing GFP- or RFP-fusion proteins, DNA fragments were PCR-amplified from genomic DNA or cDNA using primers listed in Table 1 and cloned into pDM or pCV5 vectors (Miao et al., 2017; Veltman et al., 2009) containing a multiple cloning site. To generate GxcM^4A^, the residues at position 593-596 were mutated to alanines using primers listed in Table 1. For inducible expression of constitutively active RacC, DNA fragment encoding constitutively active RacC or RFP-tagged constitutively active RacC was cloned into pDM371 or pDM359, respectively.

To generate constructs expressing GST-fusion proteins, cDNA fragment encoding Fbp17^SH3^ or the VCA domain of WASP was cloned into pGEX-6P-1 vector; cDNA fragment encoding full length Fbp17 or WASP was cloned into pFastBac1-GST vector using NdeI and XbaI sites. For expression of His-MBP-Fbp17^SH3^, cDNA fragment encoding the SH3 domain of Fbp17 was cloned into pET-MBP-3C vector using BamHI and XhoI sites.

For yeast two-hybrid assay, cDNA fragment encoding Fbp17 or the GBD domain of WASP (aa 126-230) was PCR-amplified using primers listed in Table 1 and cloned into pGADT7 prey vector. Constitutively active (CA) forms of *Dictyostelium* Rac GTPases (except for RacQ) were PCR-amplified using primers listed in Table 1 and cloned into pGBKT7 bait vector. The CA form mutations were introduced using primers listed in Table 1.

### Imaging

To image the localization of fluorescent proteins, cells were plated in coverslip chambers (Lab-Tek; NalgenNunc) filled with HL5 or LoFlo medium (Formedium) and allowed to adhere. Images were acquired on a Zeiss 880 or Zeiss 980 inverted microscope equipped with a 63 ×/1.4 oil-immersion objective. Latrunculin A treatment and chemoattractant stimulation were performed as described previously (Li et al., 2022).

For phalloidin staining, cells were plated in 8-well coverslip chamber for 2 h in HL5. Cells were fixed for 8 min at room temperature with 2% paraformaldehyde and 0.08% glutaraldehyde in KK2, permeabilized for 8 min with the addition of 0.2% TX-100, quenched in PBS containing 20 mM glycine, and washed with KK2. Cells were then stained with 14 μM Acti-stain 555 phalloidin (Cytoskeleton; cat#PHDH1-A) at room temperature in the dark for 30 min. Images were captured on a Zeiss 880 confocal microscope.

For DAPI staining, cells cultured in suspension for 60 h were gently vortexed, plated on coverslips, and allowed to adhere for 15 min. Cells were fixed with 2 ml ice-cold methanol for 5 min, washed with PBS, and then stained with DAPI containing mounting media (Abcam; cat#ab104139-20). For the experiment presented in figure S3F, cells were grown on 10 cm cell culture plate and collected for DAPI staining. Images were acquired on a Zeiss 880 confocal microscope.

Image analyses were performed using Fiji ImageJ. For experiments presented in figures 4I, 7J, S8E, S8F and S9E, the cortical-to-cytosol fluorescent intensity ratio was determined by dividing the total fluorescent intensity in the rear cortical region by that in the cytosol as described previously (Nguyen et al., 2014). For experiments presented in figures 4J and 7L, the total phalloidin intensity in the rear cortical region was measured by Fiji ImageJ.

### Migration assays

For random motility assay, vegetative cells were seeded in 6-well cell culture plate in HL5 and allowed to adhere for 4 h, except for experiment presented in figure 7C where cells were allowed to adhere overnight. Before imaging, the medium was replaced with fresh HL5. Images were acquired at 30 s intervals with phase illumination on a Zeiss 880 inverted microscope equipped with a 10×/0.45 objective. To measure the effect of GxcM overexpression on random migration, cells expressing GxcM-GFP or GFP were seeded in 8-well coverslip chamber and allowed to adhere for 60 min. Time-lapse images were collected at 30 s intervals with a Zeiss 880 microscopy equipped with a 40 ×/0.95 oil-immersion objective.

Under-agarose folic acid chemotaxis assay was performed as described before (Woznica and Knecht, 2006; Yang et al., 2021). Briefly, 5 ml 0.5% SeaKem GTG agarose melted in LoFlo medium was poured into a 50 mm glass-bottom dish (MatTek Corp). After setting of the agarose, two troughs were cut; one was filled with 1 mM folic acid and the other vegetative cell resuspended in LoFlo. Cells were allowed to migrate for 5-9 h. For experiments presented in figures 4F, 7F, S2E, and S8F, 5 ml 2% agarose was used. Images were acquired at 20 s intervals with a 10 ×/0.45 phase objective or 3 s intervals with a 63 ×/1.4 oil-immersion objective on a Zeiss 880 microscope.

To quantify migration parameters, including accumulated distance, Euclidean distance, velocity, directness and forward migration index, cells were tracked using manual tracking plugin of Fiji ImageJ (https://fiji.sc/) and analyzed using Ibidi chemotaxis tool software.

### Micropipette aspiration assay

Micropipette aspiration assay was carried out as described previously with minor modifications (Litschko et al., 2019). Briefly, a hand-made chamber was constructed using two pieces of hydrophobic glass with a space height approximately 3 mm. The chamber was filled with PBS buffer and mounted on the stage of a Nikon Eclipse Ti inverted microscope equipped with a 40×/0.75 air objective. 10 μl of cell suspension was injected into the chamber. A bovine serum albumin-coated glass micropipette with an inner diameter of 4.35 ± 0.65 μm was filled with water and positioned into the measurement chamber using a micromanipulator. Aspiration pressure was applied with a height-adjustable water reservoir. The reference pressure (0 Pa) was calibrated by observing the motion of a non-adherent cell in the micropipette. After setting the pressure difference to 500 Pa, cells were aspirated for 5 min, and a snapshot was captured at the end. Aspiration length (Lp) was determined by Fiji ImageJ.

### Protein purification

*Escherichia coli* BL21 cells transformed with GST-Fbp17^SH3^ or GST-WASP^VCA^ were grown until absorbance at 600 nm of 0.8 and induced with 0.4 mM Isopropyl β-D-1-thiogalactopyranoside (IPTG) for 16-18 h at 20°C. Bacteria pellet was resuspended in suspension buffer (50 mM Tris pH 8.0, 500 mM NaCl, 1 mM EDTA, 1 mM DTT) and lysed by sonication. Cell suspension was centrifuged at 15,000 g for 30 min to pellet the debris. The supernatant was incubated with prewashed glutathione sepharose beads (GE Healthcare) for 1-2 h at 4°C. Beads were washed three time with suspension buffer and once with washing buffer (50 mM Tris pH 8.0, 150 mM NaCl, 1 mM EDTA, 1 mM DTT). GST-fusion proteins were eluted with elution buffer (50 mM Tris pH 8.0, 10 mM reduced glutathione) followed by buffer exchange into G buffer (50 mM Tris pH 8.0, 150 mM NaCl, 1 mM EDTA) using desalting column (GE Healthcare; cat#17085101). To remove the GST tag, 10 μl PreScission Protease was added into the protein-bead mixture and incubated at 4°C overnight with gentle rotation. Proteins with the GST tag cleaved was collected from the eluate.

GST-Fbp17 and GST-WASP were purified from Sf9 insect cells. Baculovirus packaging was performed according to manufacturer’s instructions and previous description (Feng et al., 2020). Briefly, pFastBac1-GST-Fbp17 or pFastBac1-GST-WASP were transformed into DH10Bac cells to get recombinant Bacmid, which was then transformed into Sf9 cells using Cellfectin II reagent (Invitrogen; cat#10362100) to produce virus stock. The virus was then amplified by infecting Sf9 cells. After three round of infection, GST recombinant proteins were purified following the procedures described above.

His-MBP-Fbp17^SH3^ was precipitated by amylose beads (NEB; cat#E8021L). After purification, beads were preserved in buffer (25 mM Tris pH 7.5, 150 mM NaCl, 5 mM MgCl_2_, 1 mM DTT, 50% Glycerol) at -20°C.

### In vitro actin polymerization assay

Actin nucleation assay was performed as described previously with minor modifications (Diao et al., 2018) Briefly, 3 μM actin (10% pyrene labeled), 20 nM bovine brain Arp2/3 complex (Cytoskeleton; cat#RP01P), and 20 nM GST-WASP were mixed with purified Fbp17^SH3^, GST-FBP17^SH3^, or GST-Fbp17 in G buffer and incubated at room temperature. The reaction volume was 135 μl. To initiate actin polymerization, 15 μl 10 × KMEI buffer (500 mM KCl, 10 mM MgCl_2_, 10 mM EGTA, and 100 mM imidazole-HCl pH 7.4) was added. Actin assembly was traced by monitoring pyrene fluorescence by a Quanta Master Luminescence QM 3 PH fluorometer (Photo Technology International) with the excitation and emission wavelength set at 365 nm and 407 nm, respectively. 20 nM GST-WASP^VCA^ was included instead of Fbp17 proteins in one of the reactions as a positive control.

### Immunoprecipitation experiment

To identify proteins that interact specifically with GxcM, immunoprecipitation experiment and mass spectrometry analysis were carried out as described previously with minor modification (Tu et al., 2022). Briefly, cells expressing GxcM-GFP, GxcM^N790-^GFP, or Teep1-GFP were starved in DB for 3 h, lysed in ice-cold lysis buffer (10 mM Hepes pH 7.2, 100 mM NaCl, 0.5% Nonidet P-40, 10% glycerol, 1 mM NaF, 0.5 mM Na_3_VO_4_, complete EDTA-free protease inhibitor (Roche), 1 mM DTT) with the addition of 15 mM EDTA, and incubated for 5 min on ice. Lysates were centrifuged at 22,000 × g for 5 min at 4 °C. The supernatants were incubated with anti-GFP affinity beads (Smart Lifesciences; cat#SA070005) for 2 h at 4°C. Beads-bound proteins were eluted with SDS loading buffer and subjected to SDS-PAGE. Protein bands were visualized by CBB staining and subjected to in-gel trypsin digestion and mass spectrometry analysis.

For most co-immunoprecipitation experiments, cells expressing GFP- or RFP-fusion proteins were starved in DB for 3 h, lysed in ice-cold lysis buffer and incubated on ice for 5 min. Lysates were centrifuged at 22,000 × g for 5 min at 4°C. The supernatants were incubated with anti-GFP affinity beads or anti-RFP affinity beads (Smart Lifesciences; cat#SA072005) for 1 h at 4°C. Beads were washed four times in lysis buffer without protease inhibitor. Samples were eluted with SDS loading buffer and subjected to SDS-PAGE. For the experiment presented in figure 3C, cells expressing GxcM-GFP and cells expressing RFP-tagged full-length or truncations of Fbp17 were lysed separately in lysis buffer. Equal volume of lysates containing GxcM or Fbp17 were mixed and then incubated with anti-RFP affinity beads for 1 h at 4°C. Beads were washed with lysis buffer and processed for SDS-PAGE.

To verify the interaction between GxcM and RacC, cells expressing GxcM-RFP and GFP, GFP-Rac1A, or GFP-RacC were lysed in ice-cold Triton lysis buffer (10 mM Hepes pH 7.2, 100 mM NaCl, 0.1% Triton X-100, 10% glycerol, 1 mM NaF, 0.5 mM Na_3_VO_4_, complete EDTA-free protease inhibitor, 1 mM DTT) with or without 15 mM EDTA. Cleared lysates were incubated with anti-RFP affinity beads for 45 min at 4°C. Beads were washed four times with lysis buffer and processed for SDS-PAGE.

Western blotting was performed as described before (Cai et al., 2010). Anti-GFP antibody (WB, 1:5000) was purchased from Roche (cat#11814460001). Anti-DsRed antibody (WB, 1:1000), which was used to detect RFP-fusion proteins, was purchased from TaKaRa (cat#632496).

### Pull-down assays

For experiments presented in figures 3D, 3F and 5C, cells expressing GFP-fusion proteins were starved, lysed at 5×10^7^ cells/ml in lysis buffer, and incubated for 5 min on ice. Lysates were centrifuged at 22,000 × g for 5 min. Supernatants were incubated with beads containing 30 μg purified GST- or MBP-Fbp17^SH3^ for 1 h at 4°C with gentle agitation. After incubation, beads were washed four times with lysis buffer. Samples were eluted with SDS loading buffer and subjected to SDS-PAGE.

### Yeast two hybrid (Y2H) assay

To screen Rac GTPases that interact with Fbp17 or WASP, yeast two-hybrid analyses were performed using the Matchmaker GAL4 Two-Hybrid System 3 (Clontech Laboratories). *S. cerevisiae* strain AH109 was cotransfected with both bait and prey plasmids and grown on double-dropout (DD, lacking leucine and tryptophan) agar plates following manufacturer’s instructions. Clones were collected, resuspended in 100 μl H_2_O, and spotted on quadruple-dropout (QD, deficiency in leucine, tryptophan, histidine, and adenine) agar plates. The interactions between tested proteins were analyzed according to the yeast growth on QD agar plates.

### Statistics analyses

Statistical analysis was performed using GraphPad Prism. SuperPlots were generated according to (Lord et al., 2020). Statistical significance was determined by two-tailed paired or unpaired t test. In all figures, *** indicates p < 0.001, ** p < 0.01, * p < 0.05, and ns not significant.

## Acknowledgements

We thank Drs. Hong Zhang (Institute of Biophysics, CAS, Beijing, China), Junjie Hu (Institute of Biophysics, CAS, Beijing, China), Peter Devreotes (Johns Hopkins University, Baltimore, USA), and Robert Kay (MRC Laboratory of Molecular Biology, London, UK) for reagents and cells; Drs. Yanruo Zhang and Jianhui Xiao (Institute of Biophysics, CAS, Beijing, China) for helping with MPA; the proteomics core facilities in Peking University and Tsinghua University for MS analysis; the Center for Biological Imaging at the Institute of Biophysics for assistance with imaging data collection.

## Author contributions

Conceptualization, H.C. and D.L.; Methodology, H.C., D.L., S.H., and P.G.; Investigation, D.L., Y.Y., Y.W., X.C., J.H., S.S., and C.Z.; Resources, J.L.; Writing, H.C. and D.L.; Funding Acquisition, H.C. and Y.Y.; Supervision, H.C.

