## Supplemental Figures for "The GxcM-Fbp17/RacC-WASP signaling cascade regulates polarized cortex assembly in migrating cells"

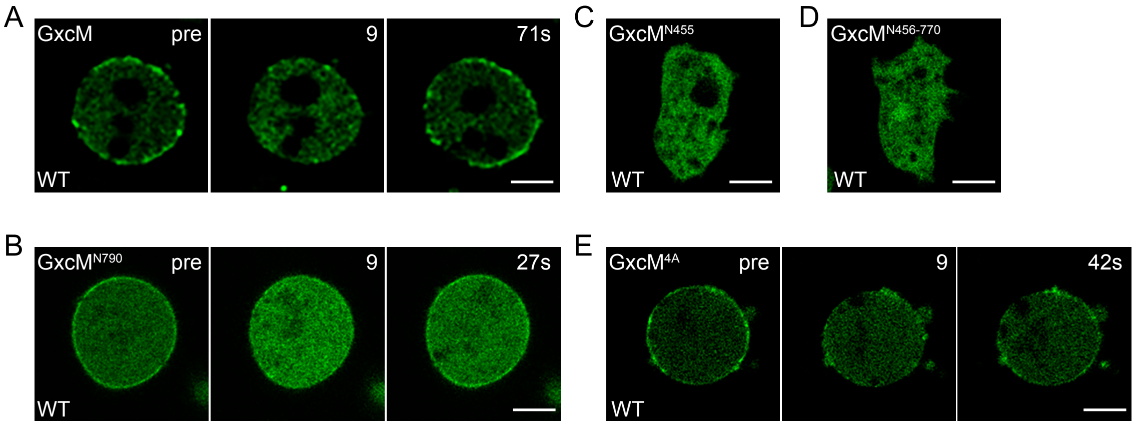


F

```
ECT2-DH  ARWQVAKELYQTESNYVNILATIIQLFQVPLEEEGQRGGPILAPEEIKTIFG 52
GxcM-DH  -RNMVINEIINTEKDYIADLNIIVNFILTPLKES-----KIIPDKDISIIFS 46

ECT2-DH  SIPDIFDVHTKIKDDLEDLIVNWDESKSIGDIFLKYSKDLVKTYPPFVNFFE 104
GxcM-DH  NIQLLLNVNKELLSDLLQKAPDEKPNHTVGTCFIYF-FKYLKIYSGYCANQT 97

ECT2-DH  MSKETIIKCEKQKPRFHAFLKINQAKPECGRQSLVELLIRPVQRLPSVALLL 156
GxcM-DH  ISADHIARCSKKIPSFKQFLEEKQASTECRQCNLESFLIKPVQRLCKYPLLL 149

ECT2-DH  NDLKKHTADENPDKSTLEKAIGSLKEVMTHINED------------------ 190
GxcM-DH  RELIKNSCEGHPDLANLEKAYTEIQEVVLSVNESKRKAEIHQKMYKIHEKLQ 201
```

Figure S1

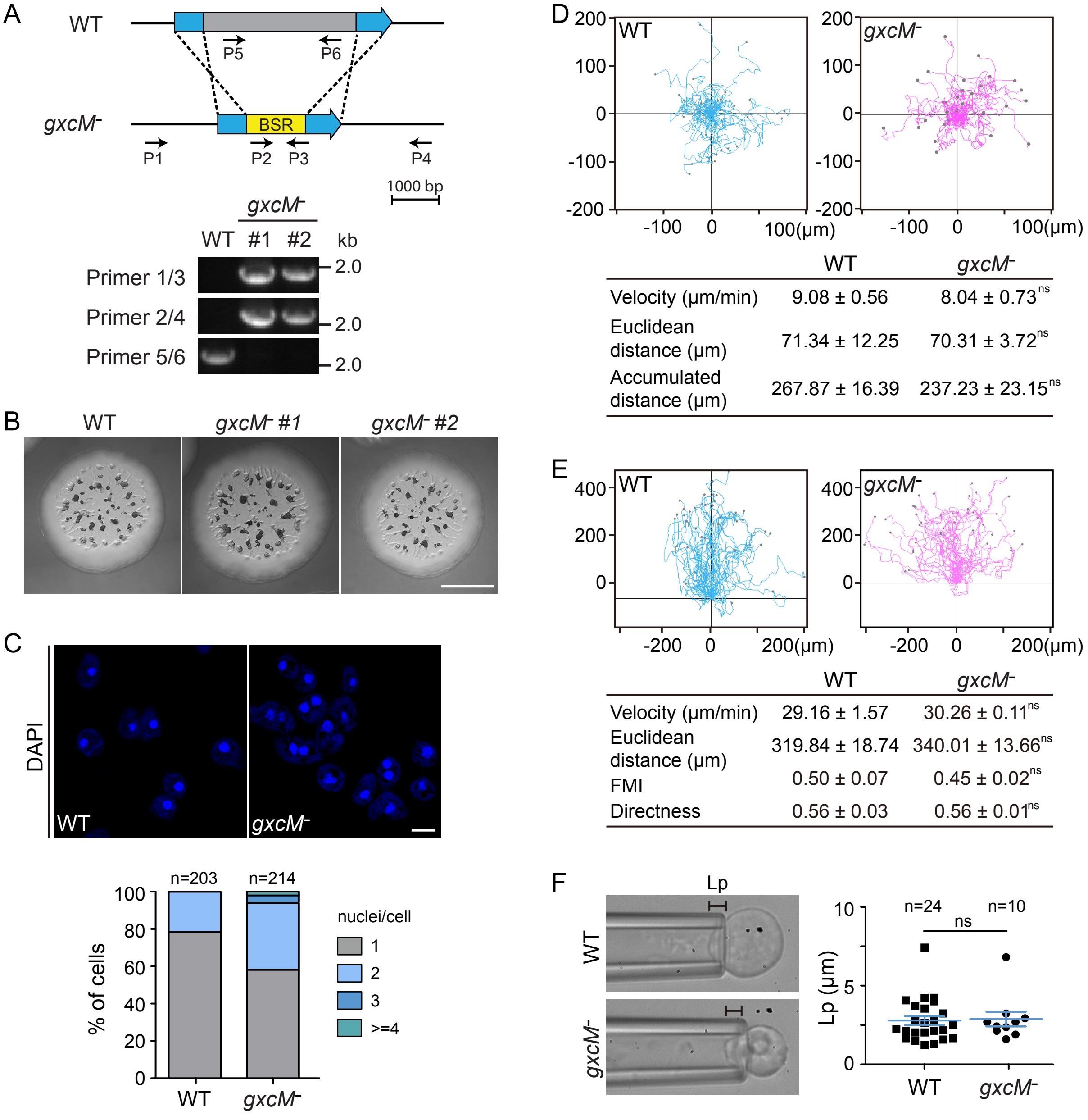


| | WT | gxcM- |
|---|---|---|
| Velocity (μm/min) | 9.08 ± 0.56 | 8.04 ± 0.73 ns |
| Euclidean distance (μm) | 71.34 ± 12.25 | 70.31 ± 3.72 ns |
| Accumulated distance (μm) | 267.87 ± 16.39 | 237.23 ± 23.15 ns |

| | WT | gxcM- |
|---|---|---|
| Velocity (μm/min) | 29.16 ± 1.57 | 30.26 ± 0.11 ns |
| Euclidean distance (μm) | 319.84 ± 18.74 | 340.01 ± 13.66 ns |
| FMI | 0.50 ± 0.07 | 0.45 ± 0.02 ns |
| Directness | 0.56 ± 0.03 | 0.56 ± 0.01 ns |

Figure S2

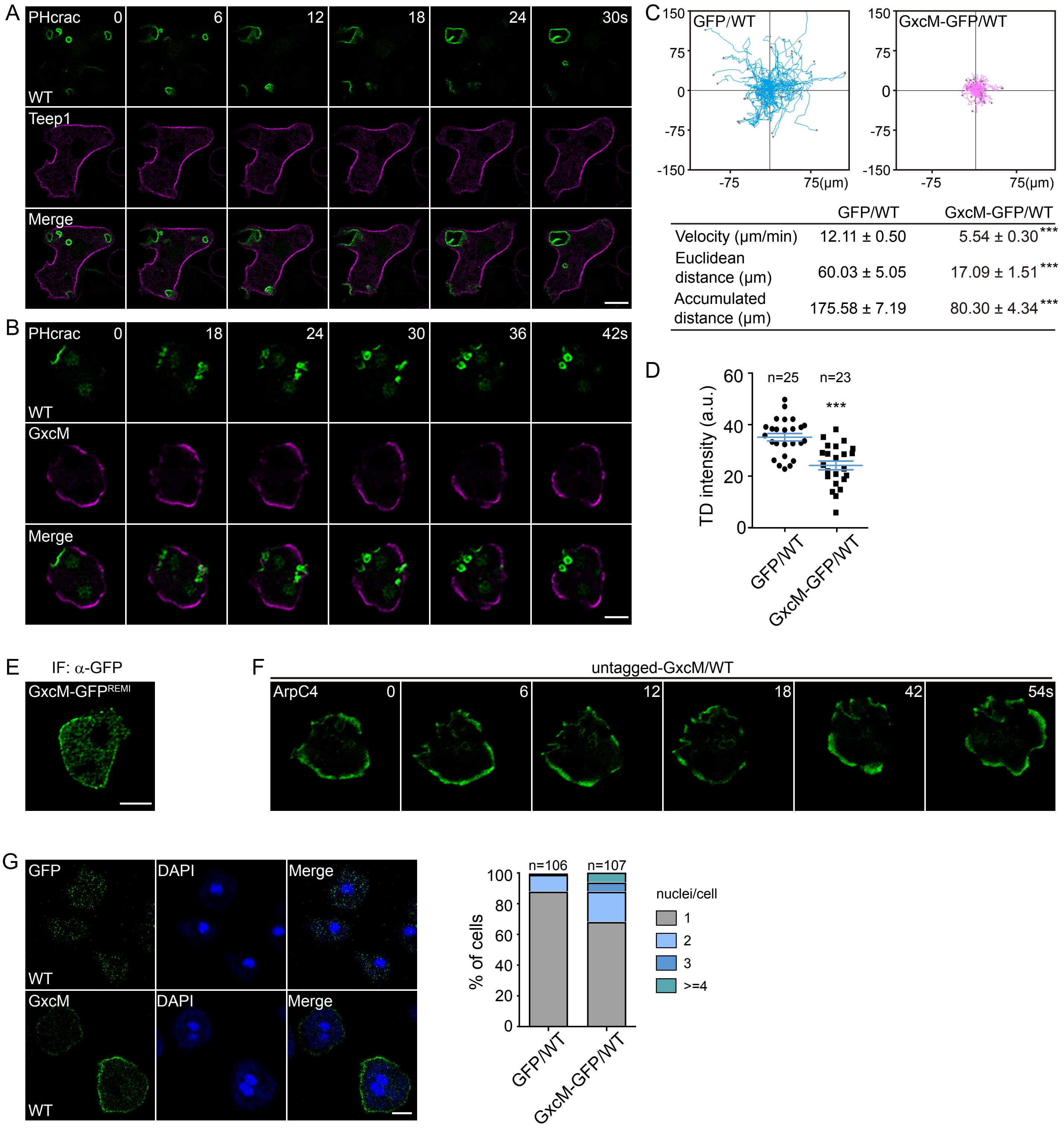
A
PHcrac
0
6
12
18
24
30s
WT
Teep1
Merge
B
PHcrac
0
18
24
30
36
42s
WT
GxcM
Merge
C
GFP/WT
GxcM-GFP/WT
150
75
0
-75
-150
-75
75(μm)
GFP/WT GxcM-GFP/WT
Velocity (μm/min) 12.11 ± 0.50 5.54 ± 0.30***
Euclidean distance (μm) 60.03 ± 5.05 17.09 ± 1.51***
Accumulated distance (μm) 175.58 ± 7.19 80.30 ± 4.34***
D
TD intensity (a.u.)
n=25
n=23
***
60
40
20
0
GFP/WT
GxcM-GFP/WT
E
IF: α-GFP
GxcM-GFP^REMI
F
untagged-GxcM/WT
ArpC4
0
6
12
18
42
54s
G
GFP
DAPI
Merge
WT
GxcM
DAPI
Merge
WT
% of cells
n=106
n=107
100
80
60
40
20
0
nuclei/cell
1
2
3
>=4
GFP/WT
GxcM-GFP/WT

Figure S3

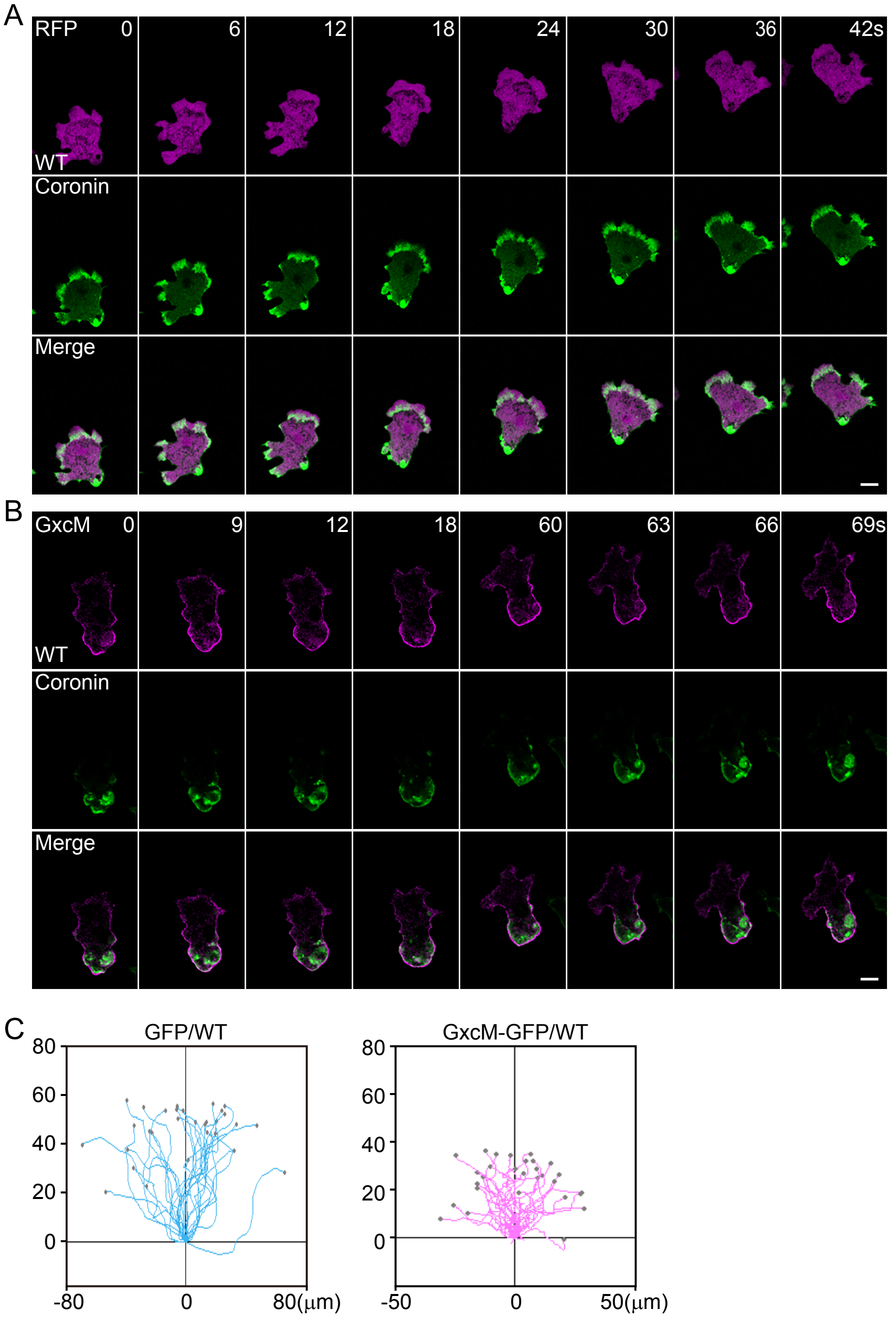


| | GFP/WT | GxcM-GFP/WT |
|---|---|---|
| Velocity (μm/min) | 32.40 ± 0.74 | 18.36 ± 0.59$^{***}$ |
| Euclidean distance (μm) | 56.03 ± 1.76 | 30.60 ± 0.94$^{***}$ |
| FMI | 0.67 ± 0.03 | 0.58 ± 0.03$^{ns}$ |
| Directness | 0.80 ± 0.02 | 0.73 ± 0.02$^{**}$ |

Figure S4

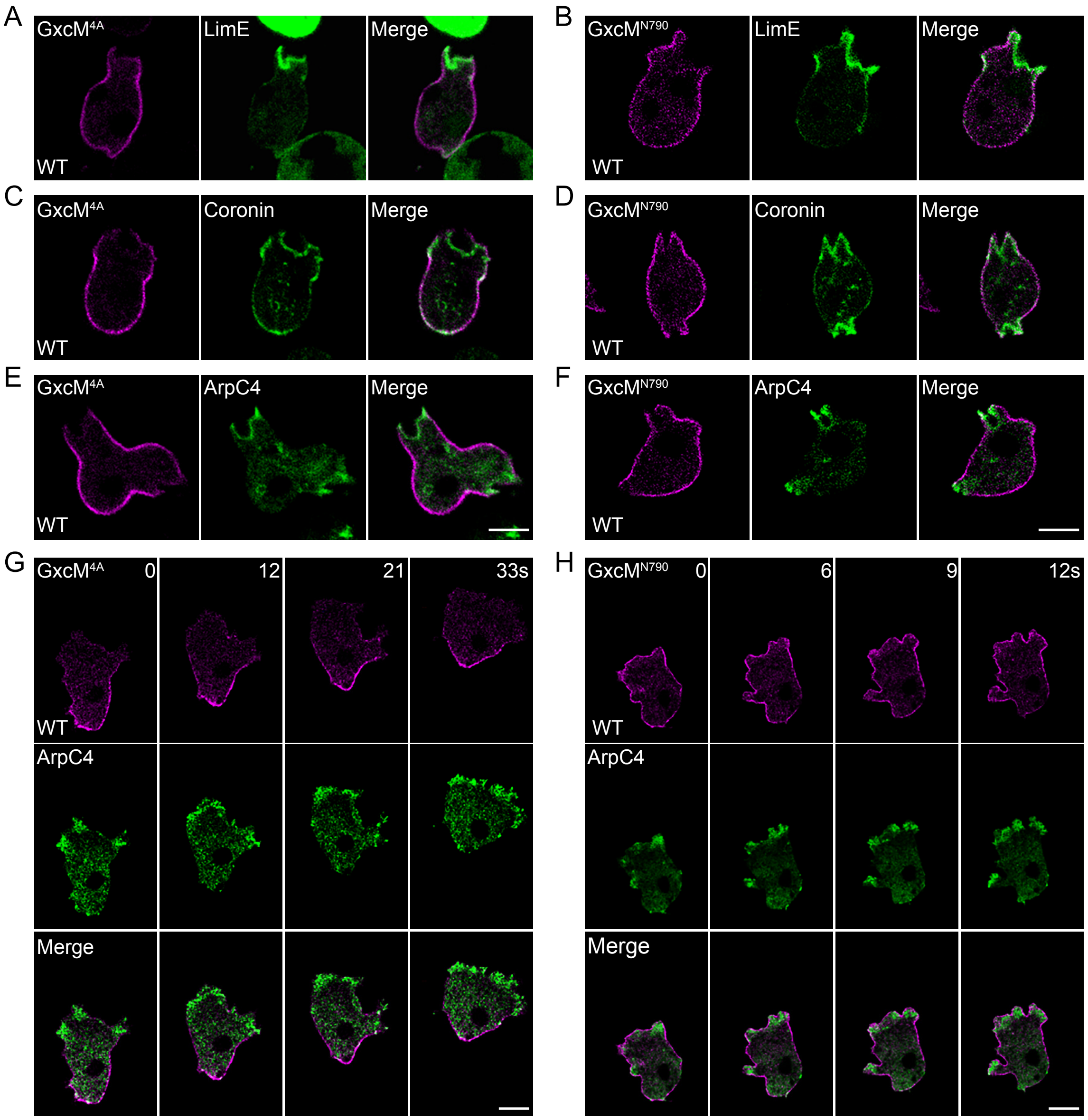
A
GxcM4A
LimE
Merge
WT
B
GxcMN790
LimE
Merge
WT
C
GxcM4A
Coronin
Merge
WT
D
GxcMN790
Coronin
Merge
WT
E
GxcM4A
ArpC4
Merge
WT
F
GxcMN790
ArpC4
Merge
WT
G
GxcM4A
0
12
21
33s
WT
ArpC4
Merge
H
GxcMN790
0
6
9
12s
WT
ArpC4
Merge

Figure S5

A

GxcM$^{791-1145}$ (SVM_score > 0.1)

791 TIATTNPSSNGISSDSPKSLHRTQTAAGALQQQQQQ

827 SKPVHAIPSKPLPPPPPKSKSSATITPTTTTTNTTV

863 VIDEPPTIVDSASTPLPSQTPIAKPRPTSTIKKTFI

899 TNSNQTSSPQQQQPATTEQAPPIPSRQQRPLSVTPT

935 TTPTPSQPVITQQNENNNTVPTKTTRPLPKPGSFTA

971 PAPVSTLINNNNTSNSSTTTTTTTTTNQPAKPIAFP

1007 SKLSNLKQAAPNNSARSASTFVPSTSHSSFVPSSVK

1043 NNTNNNNITKDDLSLPSSANNNNNNSNTDFTTPPTG

1079 KPRVMPLPPRGGSAVNVAVNNNNNINNNSPVPFPPT

1115 TNTTTPTATPPHSPQTSDLNLTKKPIIPPRK

B

| Gene ID | Accession number | Mw | #PSMs GxcM IP | #PSMs GxcM$^{N790}$ IP | #PSMs Teep1 IP |
|---|---|---|---|---|---|
| DDB_G0271812 | Q55AN3 | 52.5 KDa | 88 | 1 | 0 |

C

| Sequence (Fbp17) | # PSMs |
|---|---|
| EIPPVVQHAFEGIEK | 11 |
| KLISNPFLDGSLKESWK | 4 |
| ISLNELEQVGNQHLIFSNLNNDLSTGIDK | 3 |
| TGVTVPSHIEYQGFEGEAASGNNNSTSSPTNFR | 1 |
| EFQTFEEDRIVQSK | 3 |
| GLYDYDATCDTELSFR | 1 |
| LISNPFLDGSLKESWK | 4 |
| LIQFYANDPKEQK | 4 |
| LIQFYANDPK | 2 |
| LSVLEADYSK | 5 |
| LITAQLEELGK | 11 |
| VKGLYDYDATCDTELSFR | 1 |
| VGTYKPPTTQIKEWGLTSK | 3 |
| EFQTFEEDR | 2 |
| EAEVAHTLLSK | 10 |
| EILNTTNIKQDEYYNTDMPTLLK | 1 |
| KLISNPFLDGSLK | 4 |
| KLSVLEADYSK | 4 |
| LEQQHQEISQSIR | 3 |
| AEGELAEADKK | 1 |
| AQQADMEYR | 1 |
| AQQADMEYREILNTTNIK | 1 |
| EILNTTNIKQDEYYNTDMPTLLK | 1 |
| QWVSENR | 1 |
| LAIFVK | 6 |
| EILNTTNIK | 1 |

Figure S6

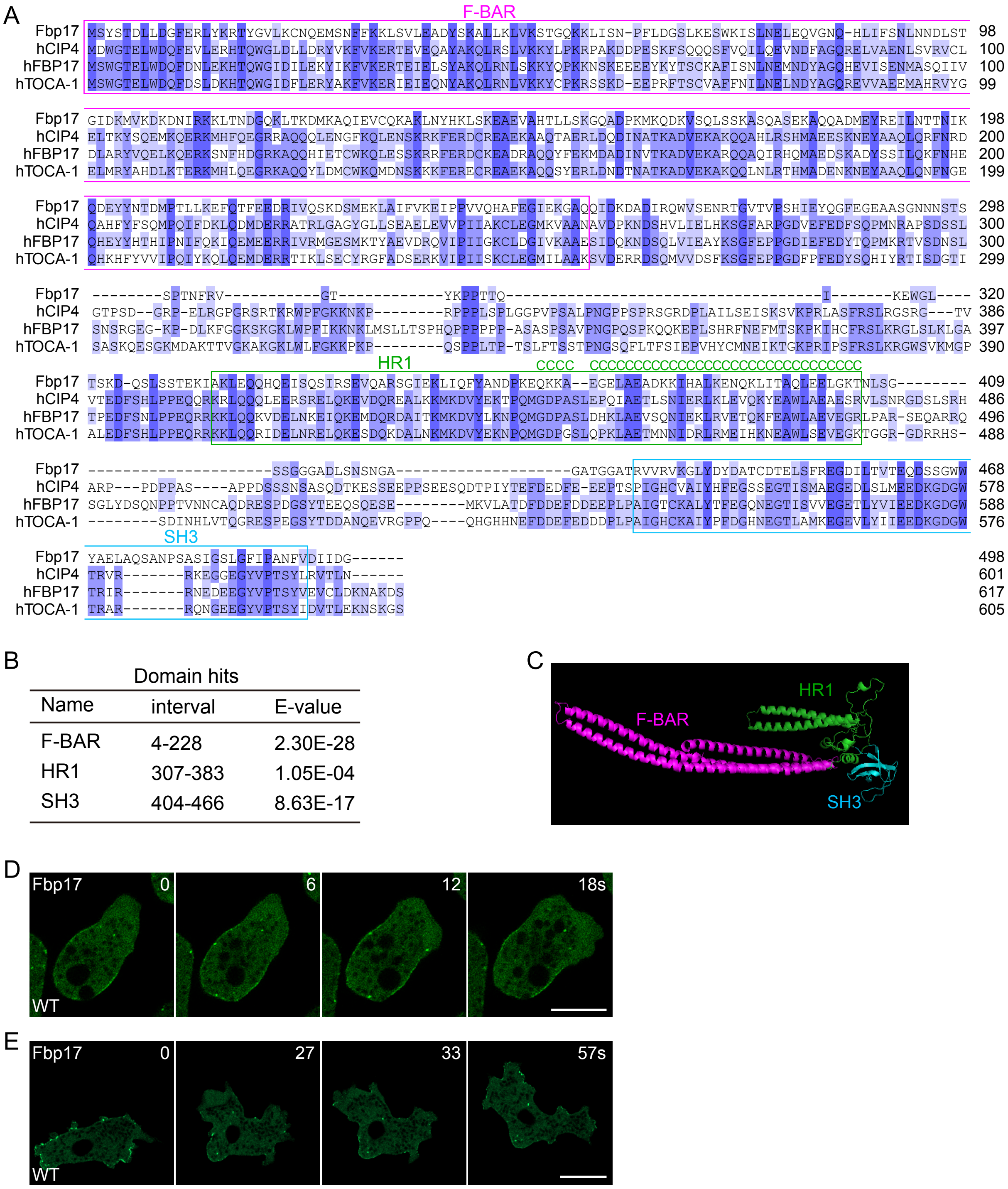


B

| Name | interval | E-value |
| --- | --- | --- |
| F-BAR | 4-228 | 2.30E-28 |
| HR1 | 307-383 | 1.05E-04 |
| SH3 | 404-466 | 8.63E-17 |

Domain hits

Figure S7

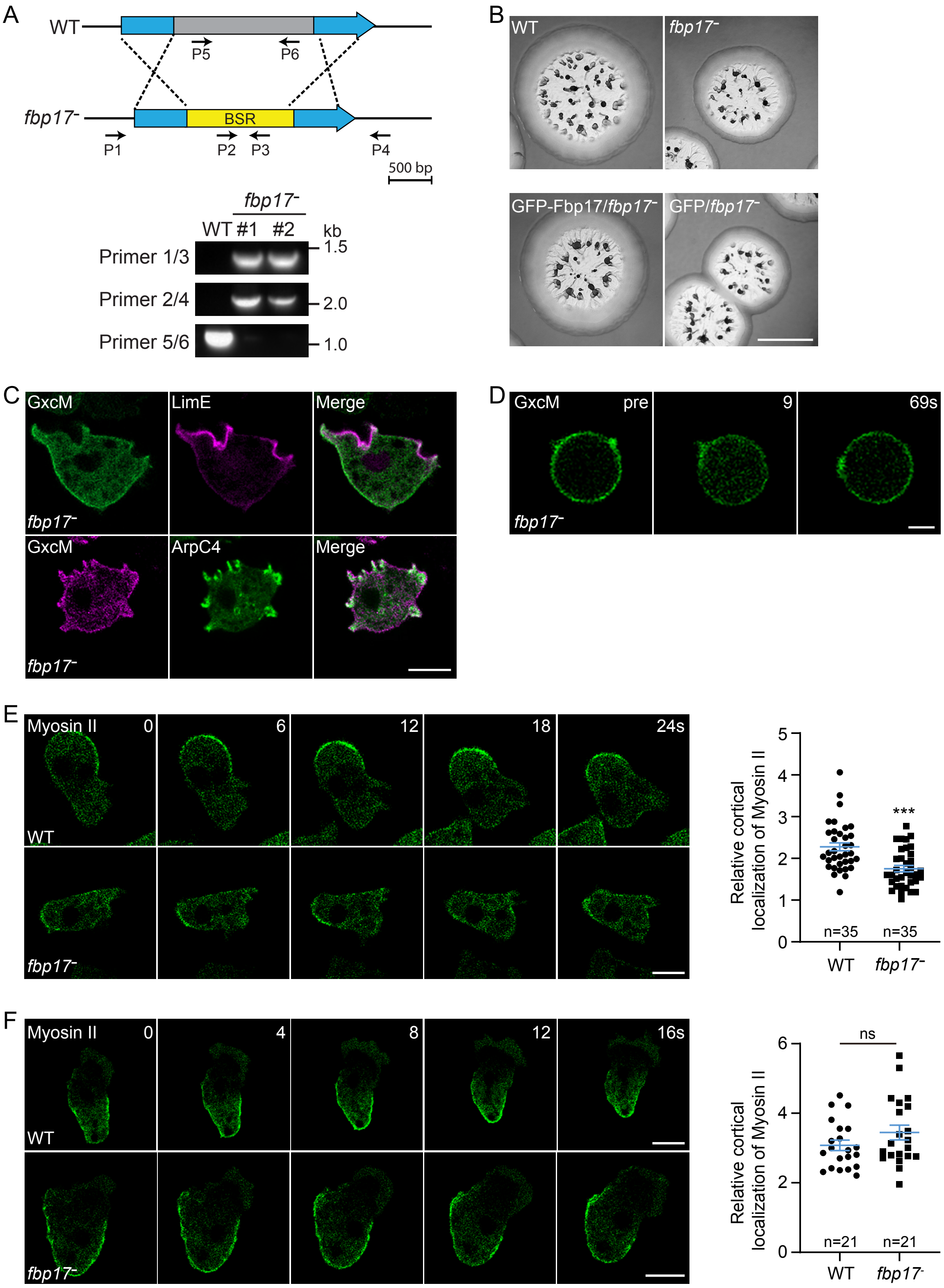

A
WT
P5
P6
fbp17⁻
BSR
P1
P2
P3
P4
500 bp
fbp17⁻
WT #1 #2
kb
Primer 1/3
1.5
Primer 2/4
2.0
Primer 5/6
1.0
B
WT
fbp17⁻
GFP-Fbp17/fbp17⁻
GFP/fbp17⁻
C
GxcM
LimE
Merge
fbp17⁻
GxcM
ArpC4
Merge
fbp17⁻
D
GxcM
pre
9
69s
fbp17⁻
E
Myosin II
0
6
12
18
24s
WT
fbp17⁻
Relative cortical localization of Myosin II
n=35
n=35
***
WT
fbp17⁻
F
Myosin II
0
4
8
12
16s
WT
fbp17⁻
Relative cortical localization of Myosin II
ns
n=21
n=21
WT
fbp17⁻


Figure S8

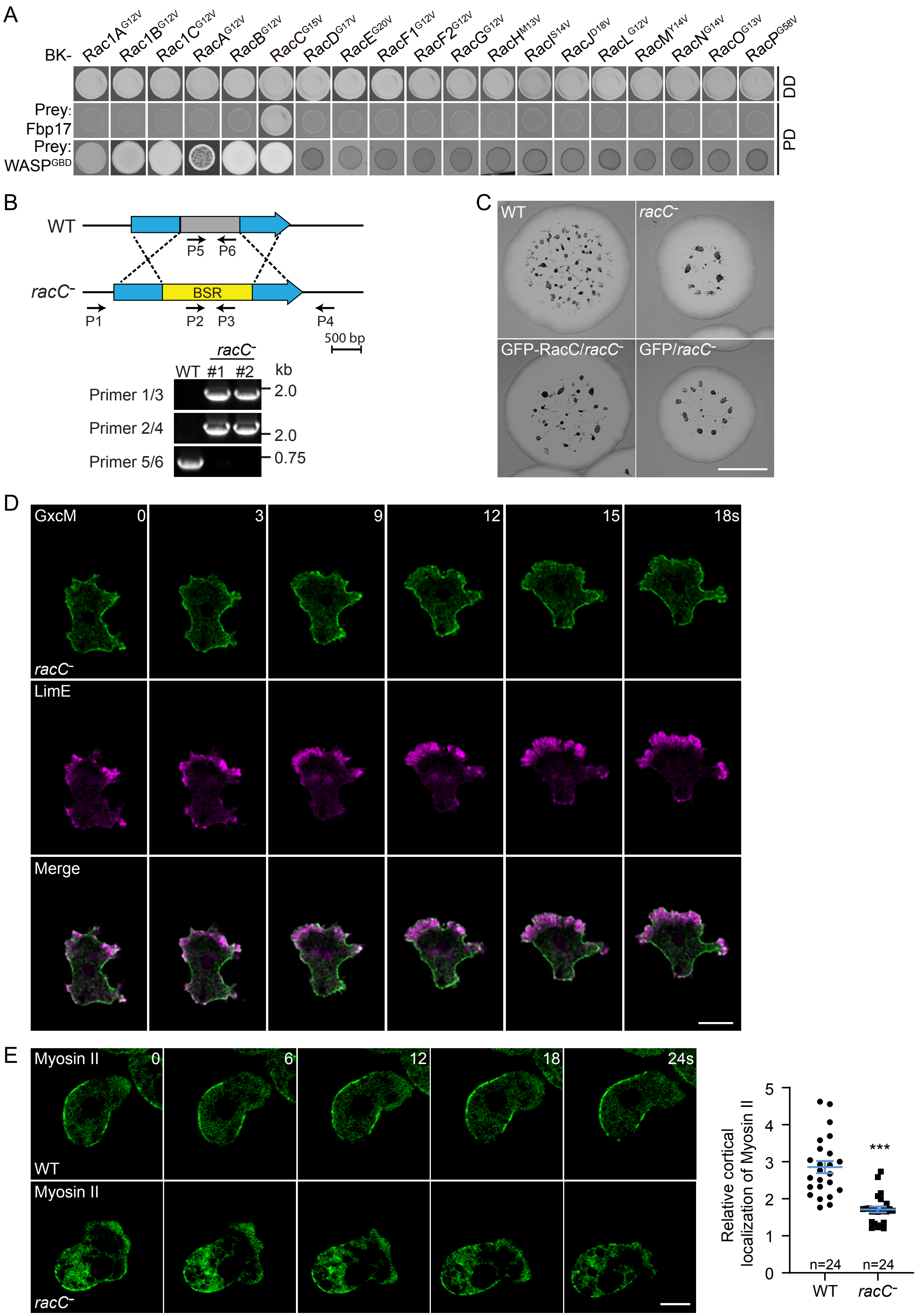
A
BK-
Rac1A$^{G12V}$
Rac1B$^{G12V}$
Rac1C$^{G12V}$
RacA$^{G12V}$
RacB$^{G12V}$
RacC$^{G15V}$
RacD$^{G17V}$
RacE$^{G20V}$
RacF1$^{G12V}$
RacF2$^{G12V}$
RacG$^{G12V}$
RacH$^{M13V}$
RacI$^{S14V}$
RacJ$^{D18V}$
RacL$^{G12V}$
RacM$^{Y14V}$
RacN$^{G14V}$
RacO$^{G13V}$
RacP$^{G58V}$
DD
Prey: Fbp17
Prey: WASP$^{GBD}$
PD
B
WT
P5 P6
racC-
BSR
P1 P2 P3 P4
500 bp
racC-
WT #1 #2 kb
Primer 1/3
Primer 2/4
Primer 5/6
2.0
2.0
0.75
C
WT
racC-
GFP-RacC/racC-
GFP/racC-
D
GxcM
0 3 9 12 15 18s
racC-
LimE
Merge
E
Myosin II
0 6 12 18 24s
WT
Myosin II
racC-
Relative cortical localization of Myosin II
0 1 2 3 4 5
***
n=24 n=24
WT racC-

Figure S9

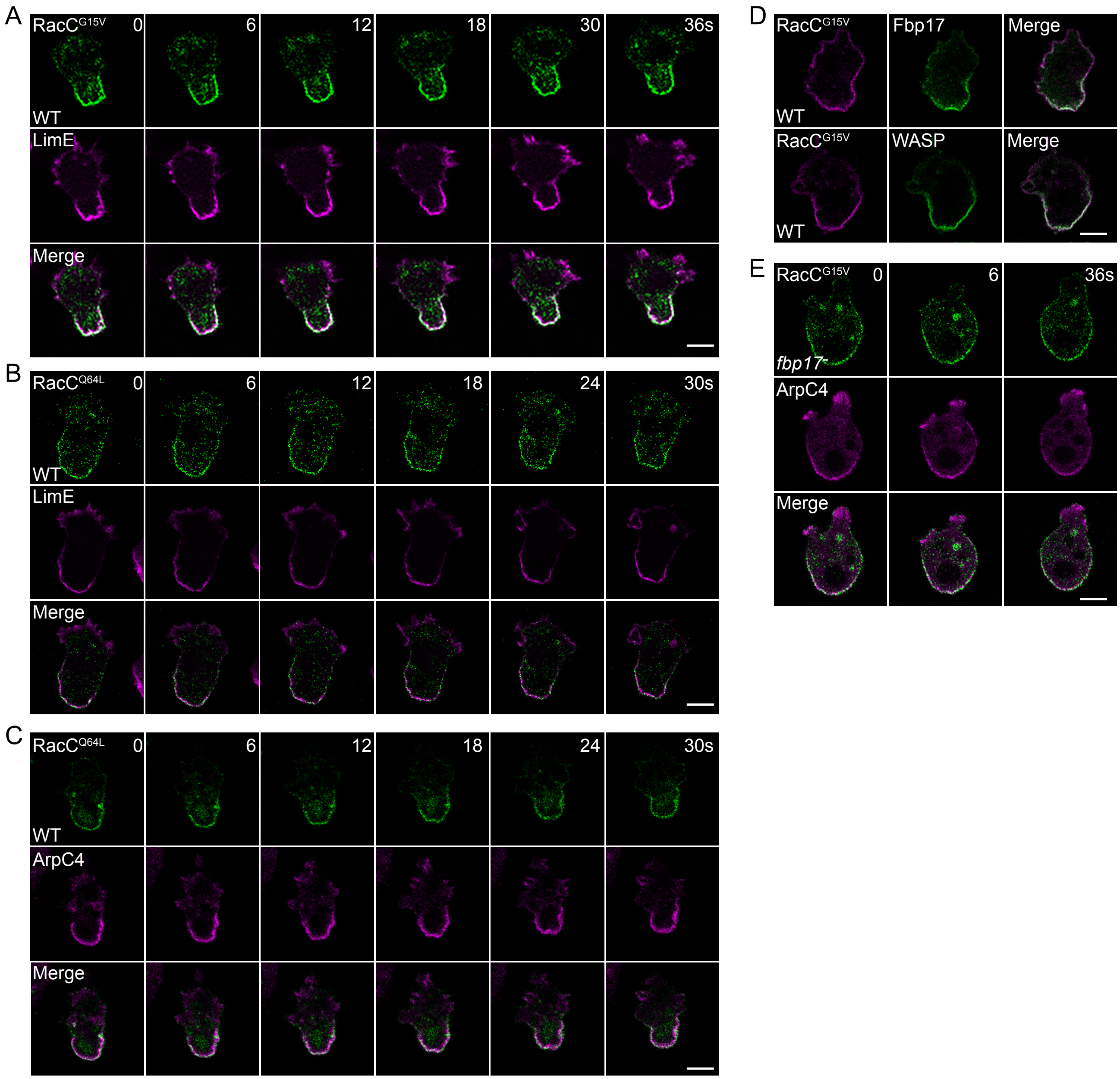
A
RacC$^{G15V}$
0
6
12
18
30
36s
WT
LimE
Merge
B
RacC$^{Q64L}$
0
6
12
18
24
30s
WT
LimE
Merge
C
RacC$^{Q64L}$
0
6
12
18
24
30s
WT
ArpC4
Merge
D
RacC$^{G15V}$
Fbp17
Merge
WT
RacC$^{G15V}$
WASP
Merge
WT
E
RacC$^{G15V}$
0
6
36s
*fbp17$^-$*
ArpC4
Merge

Figure S10

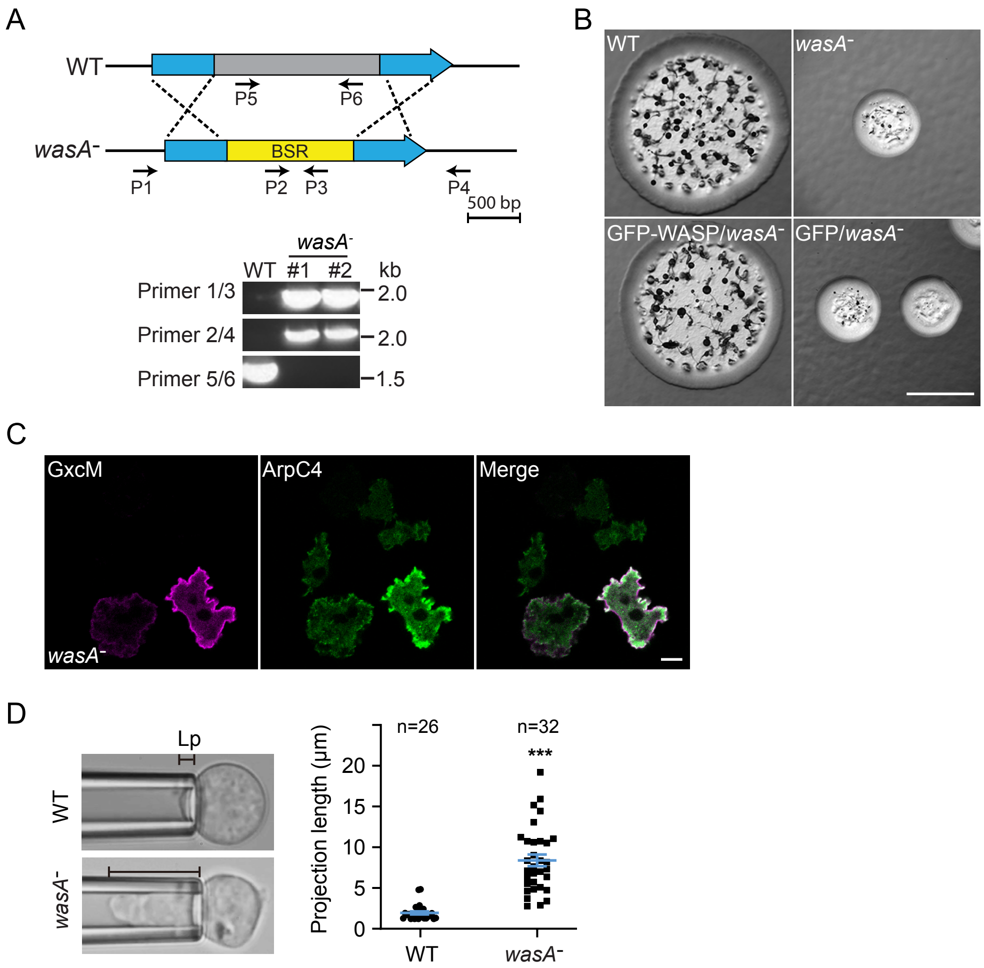
A
WT
P5
P6
wasA⁻
BSR
P1
P2
P3
P4
500 bp
wasA⁻
WT #1 #2 kb
Primer 1/3
2.0
Primer 2/4
2.0
Primer 5/6
1.5
B
WT
wasA⁻
GFP-WASP/wasA⁻
GFP/wasA⁻
C
GxcM
ArpC4
Merge
wasA⁻
D
Lp
WT
wasA⁻
n=26
n=32
***
Projection length (µm)
20
15
10
5
0
WT
wasA⁻

Figure S11

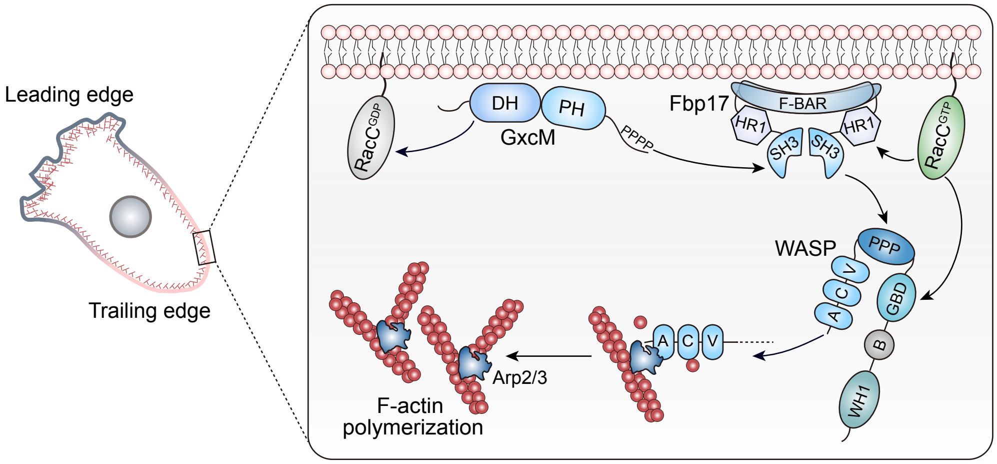
Leading edge
Trailing edge
RacC^GDP
DH
PH
GxcM
PPPP
Fbp17
F-BAR
HR1
HR1
SH3
SH3
RacC^GTP
WASP
PPP
V
C
A
GBD
B
WH1
A
C
V
Arp2/3
F-actin
polymerization

Figure S12
